# Human neural networks with sparse TDP-43 pathology reveal NPTX2 misregulation in ALS/FTLD

**DOI:** 10.1101/2021.12.08.471089

**Authors:** Marian Hruska-Plochan, Katharina M. Betz, Silvia Ronchi, Vera I. Wiersma, Zuzanna Maniecka, Eva-Maria Hock, Florent Laferriere, Sonu Sahadevan, Vanessa Hoop, Igor Delvendahl, Martina Panatta, Alexander van der Bourg, Dasa Bohaciakova, Karl Frontzek, Adriano Aguzzi, Tammaryn Lashley, Mark D. Robinson, Theofanis Karayannis, Martin Mueller, Andreas Hierlemann, Magdalini Polymenidou

**Affiliations:** Department of Quantitative Biomedicine, University of Zurich, Winterthurerstrasse 190, 8057 Zurich, Switzerland; Department of Molecular Life Sciences, University of Zurich, Winterthurerstrasse 190, 8057 Zurich, Switzerland; SIB Swiss Institute of Bioinformatics, University of Zurich, Winterthurerstrasse 190, 8057 Zurich, Switzerland; Department of Biosystems Science and Engineering, ETH Zürich, Mattenstrasse 26, 4058 Basel, Switzerland; Brain Research Institute, University of Zurich, Winterthurerstrasse 190, 8057 Zurich, Switzerland; Institute of Neuropathology, University of Zurich, Schmelzbergstrasse 12, 8091 Zurich, Switzerland; Department of Histology and Embryology, Faculty of Medicine, Masaryk University Brno, Kamenice 3, 62500, Brno, Czech Republic; Queen Square Brain Bank for Neurological diseases, Department of Movement Disorders, UCL Institute of Neurology, London, WC1N 1PJ, UK; Department of Neurodegenerative Disease, UCL Institute of Neurology, Queen Square, London, WC1N 3BG, UK

**Author notes:** These authors contributed equally to this work (Katharina M. Betz, Silvia Ronchi).

## Abstract

Human cellular models of neurodegeneration require reproducibility and longevity, which is necessary for simulating these age-dependent diseases. Such systems are particularly needed for TDP-43 proteinopathies^1,2^, which involve human-specific mechanisms^3–6^ that cannot be directly studied in animal models. To explore the emergence and consequences of TDP-43 pathologies, we generated iPSC-derived, colony morphology neural stem cells (iCoMoNSCs) via manual selection of neural precursors^7^. Single-cell transcriptomics (scRNA-seq) and comparison to independent NSCs^8^, showed that iCoMoNSCs are uniquely homogenous and self-renewing. Differentiated iCoMoNSCs formed a self-organized multicellular system consisting of synaptically connected and electrophysiologically active neurons, which matured into long-lived functional networks. Neuronal and glial maturation in iCoMoNSC-derived cultures was similar to that of cortical organoids^9^. Overexpression of wild-type TDP-43 in a minority of iCoMoNSC-derived neurons led to progressive fragmentation and aggregation, resulting in loss of function and neurotoxicity. scRNA-seq revealed a novel set of misregulated RNA targets coinciding in both TDP-43 overexpressing neurons and patient brains exhibiting loss of nuclear TDP-43. The strongest misregulated target encoded for the synaptic protein NPTX2, which was consistently misaccumulated in ALS and FTLD patient neurons with TDP-43 pathology. Our work directly links TDP-43 misregulation and NPTX2 accumulation, thereby highlighting a new pathway of neurotoxicity.

---

TAR DNA-binding protein 43 (TDP-43) accumulates in affected neurons from patients with neurodegenerative diseases, including amyotrophic lateral sclerosis (ALS) and frontotemporal lobar dementia (FTLD)^1,2^. TDP-43 is an essential RNA-binding protein^10–12^ that is tightly autoregulated via binding to its own mRNA^13–16^. At the physiological state, TDP-43 is predominantly nuclear and directly controls the processing of hundreds of its RNA targets^13,17^. Conversely, TDP-43 forms pathological aggregates in disease, which are neurotoxic *per se*, featuring a potency that correlates with disease duration in FTLD patients^18,19^. Moreover, the aggregates trap newly synthesized TDP-43, leading to nuclear clearance and loss of its normal function^1,20,21^. This effect has detrimental consequences as it leads to broad splicing misregulation^13,22^, including the inclusion of cryptic exons in specific TDP-43 RNA targets^23^, such as *STMN2^3,4^* and *UNC13A^5,6^*. Both these RNA targets are neuronal and human-specific, and their levels were recently found to be significantly reduced in brains of patients with TDP-43 proteinopathies, directly linking the loss of TDP-43 nuclear function to neurodegeneration.

The recognition of *STMN2* and *UNC13A* motivated the development of fully human experimental models for TDP-43 proteinopathies in order to decipher the key targets and downstream pathological mechanisms of TDP-43 misregulation. Induced pluripotent stem cell (iPSC)-based systems offer this possibility, and several breakthroughs were made in recent years with this technology, including the generation and characterization of numerous iPSC lines from ALS and FTLD patients^24^, the recognition of early neuronal phenotypes in human neurons with ALS-linked mutations^25,26^ and disease-linked transcriptomic signatures^27^. Yet, most studies with iPSC-derived neurons from patients with TDP-43 proteinopathies reported low to no TDP-43 pathologies^28–30^, potentially due to the early maturation state of human neurons in culture.

## iCoMoNSCs are uniquely homogeneous

We generated a self-renewing human neural stem cell line (designated iCoMoNSCs) via manual selection based on their colony morphology^7^, from induced pluripotent stem cells (iPSCs), which we derived from normal human skin fibroblasts through episomal reprogramming (Extended Data Fig. 1a). iCoMoNSCs were stable across at least 24 passages retaining their characteristic radial morphology in cell clusters and karyotype, as well as expression of NSC-specific markers (Fig. 1a, Extended Data Fig. 1b-f). To determine the homogeneity of our iCoMoNSCs, we performed single-cell RNA sequencing (scRNA-seq) of two biological replicates at passage 22. Pre-processing, quality control and filtering yielded >8300 cells per replicate with a median number of ~2000 detected genes and ~4800 unique molecular identifiers (UMIs) (Extended Data Fig. 1g,h), which were separated in tightly associated clusters (Extended Data Fig. 1i), mostly defined by cell cycle stage (Extended Data Fig. 1j) and composed of cells from both biological replicates (Extended Data Fig. 1k,l), demonstrating that our iCoMoNSCs were extremely homogeneous. The classical NSC marker genes *NES*, *SOX2*, *NR2F1* and *CDH2*, as well as IRX2 and SOX1, were expressed in a subset of cells from all clusters with similar levels (Extended Data Fig. 1m), suggesting that the majority of cells were true, self-renewing neural stem cells at different cell cycle stages. We then identified cluster marker genes (Extended Data Fig. 2a) and interrogated the expression of sets of known cell type marker genes (Extended Data Fig. 2b). This showed that most of the cells were in G1/S phase (clusters 0, 1 and 5), followed by ~19% of cells (cluster 2) marked by classical cell cycle-associated genes (G2/M). For a minority of the cells, markers and specifically expressed genes indicated lower multipotent states with either gliogenic (~13% of cells, cluster 3) or neurogenic (~8% of cells, cluster 4) nature. Finally, for a very small percentage of cells (~0.3%, cluster 6) the expression of neuron-specific genes and cluster markers demonstrated their committed neuroblast nature (Extended Data Fig. 1i,l, Extended Data Fig. 2a-b). Taken together, our data showed that up to 79% of all cells present in our iCoMoNSCs were true, self-renewing neural stem cells.

**Figure 1.**
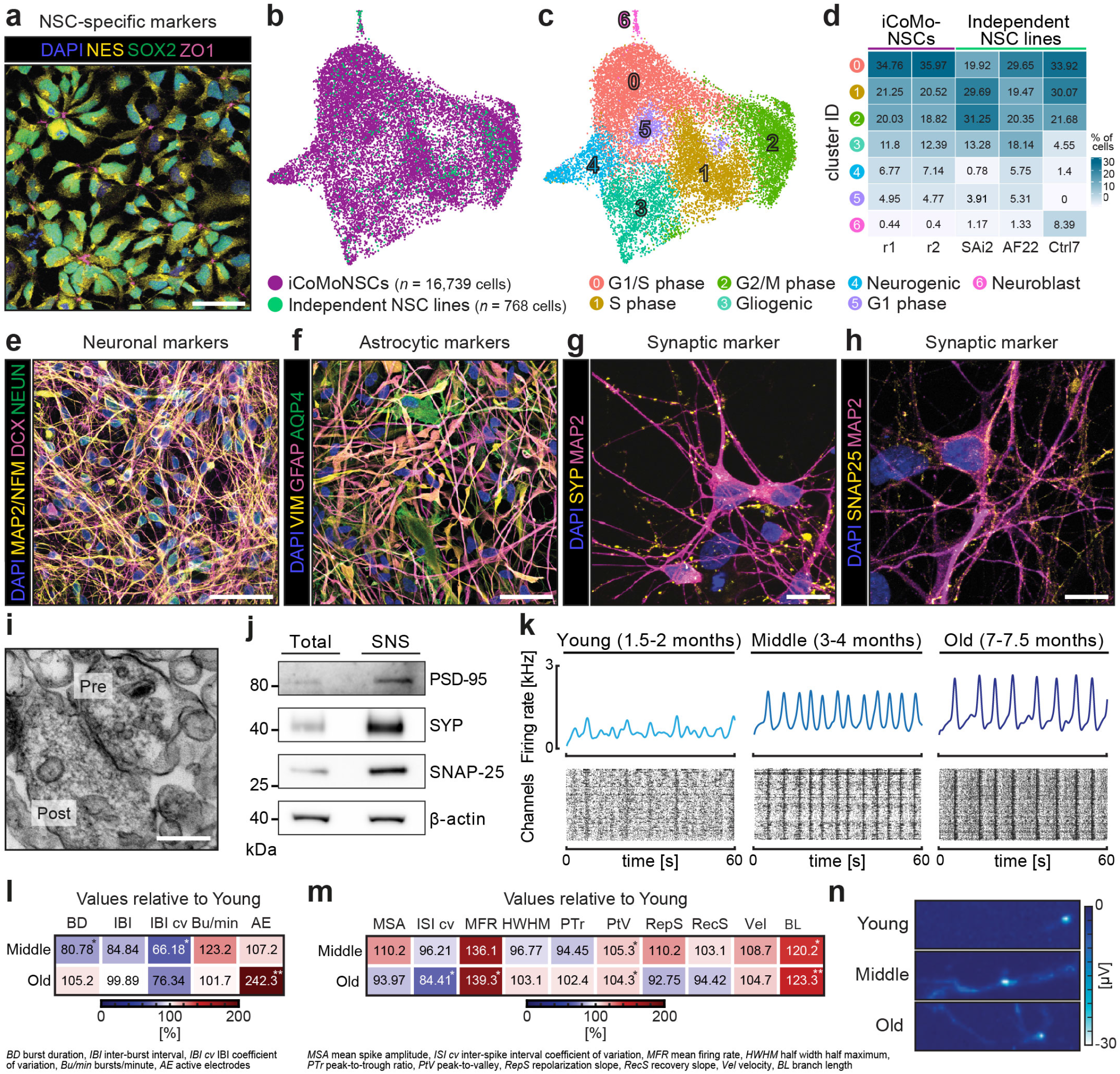
iCoMoNSC neurons form functional networks. **a**, Immunofluorescence of iCoMoNSCs for NSC markers. **b**, UMAP of iCoMoNSCs integrated with 768 cells from three independent NSC lines and (**c**) corresponding annotated clusters. **d**, Percentage of cell distribution per sample across clusters. **e**, Immunofluorescence of human neural cultures stained with neuronal or (**f**) astrocytic markers. Synaptic marker immunofluorescence at 4 months for (**g**) SYP and (**h**) SNAP-25. **i**, Electron microscopy of SNS showing pre- and postsynaptic compartments. **j,**Western blots of SNS fractions. **k**, Population spike time histograms (top) and raster plots (bottom) of neuronal networks. **l**, Heatmaps of percentage changes in network or (**m**) single-cell and subcellular metrics features. **n**, Spike-sorted axonal branches. Scale bars, (**a**,**e**,**f**) 50 μm, (**g**,**h**) 10 μm, (**i**) 250 nm.

To corroborate our results, we integrated our data with previously published scRNA-seq datasets from independent human neural stem cells (NSCs)^8^, which were distributed amongst our iCoMoNSCs in all clusters (Fig. 1b-d, Extended Data Fig. 2c). This allowed us to refine our cluster annotation (Fig. 1c). Next, we compared the cluster abundances of all samples individually and found that the iCoMoNSCs and the iPSC-derived AF22 and Ctrl7 lines had the most similar cell distributions, whereas the SAi2 line that was derived from human fetal hindbrain primary NSCs showed slightly different cell cycle distribution and approximately 9-fold less neurogenic cells (Fig. 1c,d). Despite the similarities to the independent NSC lines, our iCoMoNSCs contained significantly fewer (between 3 and 20 times) committed neuroblasts, represented in cluster 6 (Fig. 1d), indicating that our iCoMoNSCs consist primarily of self-renewing neural stem cells and significantly fewer postmitotic, committed neuroblasts than the independent NSC lines.

## iCoMoNSC neurons form functional networks

Upon differentiation (Extended Data Fig. 3a), our iCoMoNSCs consistently generated mixed neuronal and glial multilayer cultures (Fig. 1e,f)^31,32^. After 1.5 months of differentiation, these cultures consisted of approximately 30% of NEUN+neurons, a percentage that stabilised at approx. 35% at later time points (Fig. 1e, Extended Data Fig. 3b,c). In contrast, the number of Ki67+proliferating cells dropped from virtually 100% in iCoMoNSCs (Extended Data Fig. 3d) to ~5% (Extended Data Fig. 3e). To investigate the presence of synaptic markers, we first immunolabeled 3-month-old cultures for SYP (synaptic vesicles; Fig. 1g) and SNAP-25 (synaptic vesicle fusion machinery; Fig. 1h) and found a typical punctate pattern. To further assess synapse formation, we prepared synaptoneurosomes (SNS)^33,34^, a subcellular preparation, enriched in resealed presynaptic and postsynaptic structures. SNS fractions from 3-month-old human neural cultures were analyzed by transmission electron microscopy (TEM), which revealed typical synaptic morphology, consisting of both presynaptic vesicles and postsynaptic densities (Fig. 1i). Immunoblots of total lysates and SNS fractions, prepared from the human neural cultures, confirmed the enrichment of synaptic markers (Fig. 1j).

To assess the functionality of these synapses, we first performed *in vitro* two-photon calcium imaging after bolus-loading of the 3-month-old cultures with the membrane-permeable ester form of the calcium indicator Oregon Green BAPTA-1. Calcium transients of recorded neuronal somata demonstrated that the cultures indeed displayed sparse spontaneous activity patterns (Extended Data Fig. 3f). To formally confirm neuronal activity in our cultures, we assessed their electrophysiological properties via whole-cell patch-clamp measurements. 3.5-month-old patched neurons (Extended Data Fig. 3g) had a hyperpolarized resting membrane potential (−59.7 ± 4.3 mV, n = 7) and fired single or multiple action potentials upon depolarizing current injection (10 out of 11 neurons, Extended Data Fig. 3h-i). Voltage-clamp recordings showed rapidly inactivating inward currents and slowly inactivating outward currents, typical for Na^+^- and K^+^-currents, respectively (peak Na+ current density: −86.7 ± 20.5 pA/pF; peak K^+^ current density: 149.4 ± 28.0 pA/pF; n = 10; Extended Data Fig. 3j). Collectively, these analyses demonstrated that iCoMoNSC-derived neurons contained voltage-dependent channels and were electrophysiologically active with a hyperpolarized resting membrane potential.

To investigate whether iCoMoNSC-derived neurons were interconnected and displayed coordinated activity, we used high-density microelectrode arrays (Extended Data Fig. 4b; HD-MEAs)^35,36^. Neural cultures were plated onto the HD-MEAs after approximately 1, 3 and 6 months of differentiation and were then allowed to reconnect for one month before recording to compare cultures denoted in the rest of the manuscript as young (1.5-2 months), middle (3-4 months) and old (7-7.5 months) (Extended Data Fig. 4a). Young cultures exhibited a lower burst activity than middle and old (Fig. 1k). We then analyzed burst metrics (Fig. 1l, Extended Data Fig. 4c,f-j)^37^ and found a significant decrease in burst duration (BD) between young and middle cultures (*p*<0.025), as well as a decrease in the inter-burst interval coefficient of variation (IBI cv) (*p*<0.025) (Fig. 1l, Extended Data Fig. 4f,h). The increase of IBI cv in young cultures compared to middle (Fig. 1l, Extended Data Fig. 4h) suggested that early maturation stages were characterized by irregular bursts. Additionally, for 30% of the young cultures a network analysis could not be conducted as bursts were undetectable, which is indicative of still-developing synaptic connections. In contrast, older cultures showed detectable bursts in the majority of MEAs (>90%). Additionally, the percentage of active electrodes (AE) increased 2.4-fold (*p*<0.001) from young to old cultures (Fig. 1l, Extended Data Fig. 4j).

We next compared the single-cell and subcellular resolution metrics (Fig. 1m, Extended Data Fig. 4d,e,k-t)^38^. We found a 1.2-fold increase (*p*<0.025) in inter-spike interval coefficient of variation (ISI cv)^37^ between young and old cultures (Fig. 1m, Extended Data Fig. 4l), indicating more irregular firing rates at later developmental stages. In addition, a 0.7-fold lower mean firing rate (MFR) in young compared to old cultures demonstrated an increase in spontaneous activity over time (Fig.1m, Extended Data Fig. 4m). Longer-lasting action potential recovery times were evidenced by the peak-to-trough ratio metric (PtV), which increased from young to middle (*p*<0.025) and from young to old cultures (*p*<0.025) (Fig. 1m, Extended Data Fig. 4p).

Next, the subcellular resolution features branch length (BL) and action-potential-propagation velocity were used to compare the three developmental stages. Significant differences were found in the neuron BL between young and middle (*p*<0.025), and between young and old cultures (*p*<0.001), indicating different functional maturation stages (Fig. 1m, Extended Data Fig. 4t). Principal component analysis (PCA) showed a separation of all three maturation stages based on all 15 analyzed parameters (Extended Data Fig. 4u). Taken together, these functional metrics indicated increased maturation upon aging of iCoMoNSC-derived neurons in culture.

## iCoMoNSC neurons and glia resemble brain organoids

To further characterize our iCoMoNSC-derived cultures, we performed scRNA-seq of young, middle and old (Extended Data Fig. 4a), in biological duplicates. After processing we retained 3500-8500 cells per sample with 2800-4800 detected genes and 6000-16000 UMIs. Cells were distributed across 19 clusters, which self-organized with groups of clusters on the top left quadrant representing neurons, whereas the bottom left quadrant contained clusters of astrocytes and other glial cells (Fig. 2a). Neuronal and astrocytic clusters showed increasing maturation over time in differentiation, which was evident upon visualization of all experimental (Fig. 2b) or cell cycle (Fig. 2c) stages. Cluster abundance analysis (Extended Data Fig. 5a-c) and the visualization of individual, cell type-specific markers revealed that our cultures matured over time: NSC-specific markers, such as *SOX2*, *NQO1,* had high expression in iCoMoNSCs (Extended Data Fig. 5d). Astrocyte maturation was highlighted by marker *GFAP* (astrocytes), which was expressed in middle and old neural cultures, *PTPRZ1* (oligodendrocyte progenitor cells, OPCs) was detected in all samples and *DCN* (pericytes) in young and middle (Extended Data Fig. 5e). Neuronal maturation is highlighted by markers *SYP* (neuronal) and *SLC32A1* (GABAergic), which were detected in young, middle and old cultures, whereas *SLC17A6* (glutamatergic) was mostly detected in old cultures (Extended Data Fig. 5f). Clusters were manually annotated based on analysis of cluster markers (Extended Data Fig. 5b,g), known markers (Extended Data Fig. 6), CoDex (Cortical Development Expression) viewer^39^, PanglaoDB database^40^ and UCSC Cell Browser. In line with the increasing maturation over time in differentiation, young neurons were annotated as young inhibitory (cluster 2), excitatory (cluster 3) and dopaminergic neurons (cluster 13), as well as apoptotic neurons (cluster 15), while at the middle stage, their annotations converted into maturing (cluster 6) or excitatory glutamatergic (cluster 7) and in the old cultures, neuronal subtypes were clearly defined as GABAergic (cluster 5) and glutamatergic (cluster 7). Similarly, gliogenic clusters were annotated as glial precursors (cluster 18), radial glia/early astrocytes (cluster 10), radial glia (cluster 12) and glia (cluster 11) in young cultures, young astrocytes (cluster 9) and glia (cluster 11) at the middle stage, and mature astrocytes (cluster 4), astrocytes (cluster 16), and OPCs (cluster 8) in the old cultures. A small percentage of glial cells (between 0.16 – 0.7%) was identified as pericytes (cluster 17), regardless of the maturation stage (Fig. 2a-c, Extended Data Fig.4a, Extended Data Fig. 5a-f).

**Figure 2.**
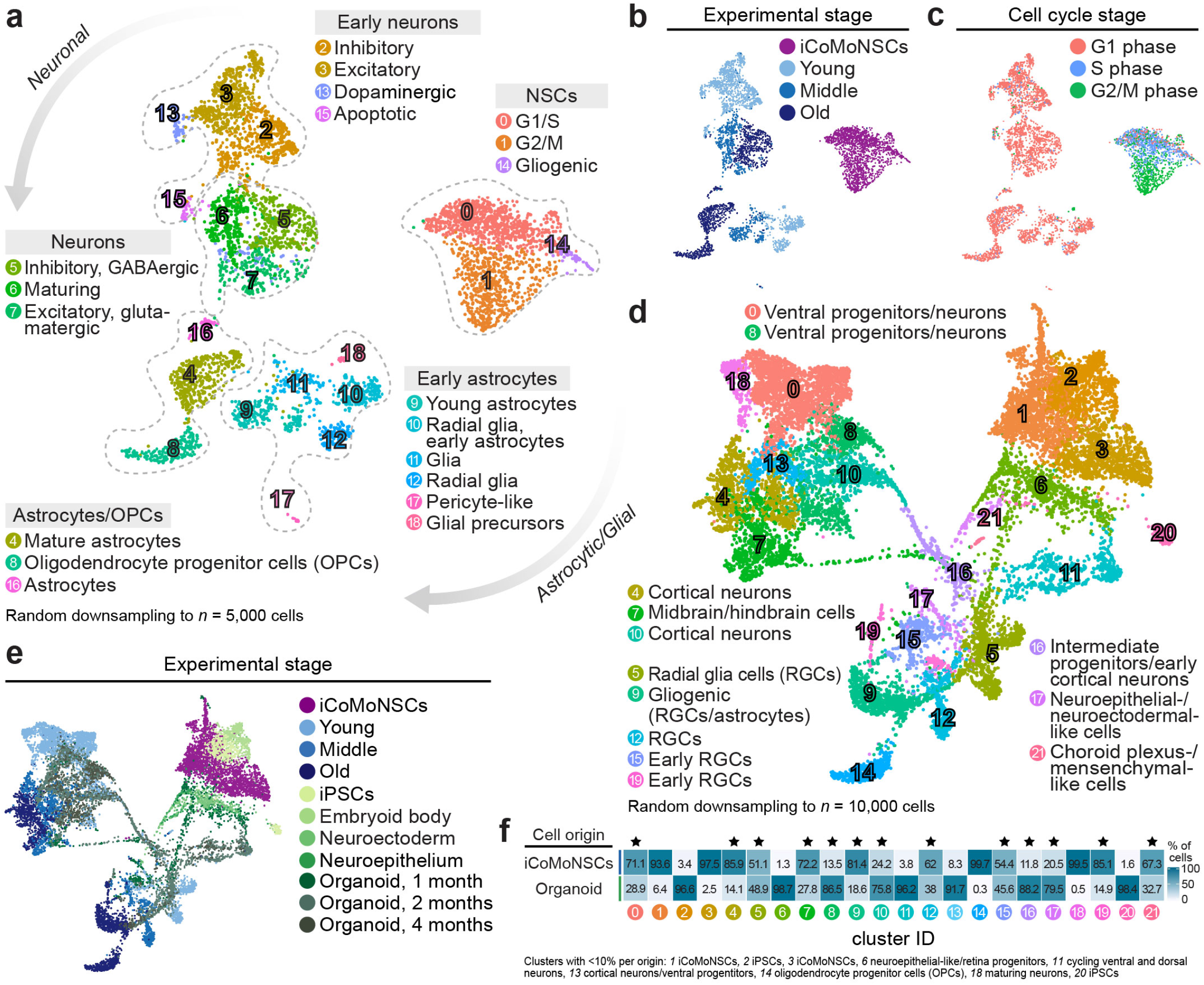
iCoMoNSC neurons and glia partially resemble brain organoids. **a**, UMAP of young, middle and old iCoMoNSC-derived neural cultures. Colors highlight manually annotated clusters, (**b**) experimental or (**c**) cell cycle stage. **d**, All samples from (**a**) integrated with organoid datasets. Annotation of clusters consisting of both iCoMoNSC and organoid cells are indicated. **e**, Same UMAP as (**d**) highlighting the experimental stage origin. **f**, Cell distribution across all clusters. Stars indicate clusters with cell composition of at least 10% per origin.

To determine the level of maturation of the emerging cell types within our iCoMoNSC-derived cultures, we integrated our data with two brain organoid scRNA-seq datasets^9^ (Fig. 2e, Extended Data Fig. 5h) and identified 22 clusters (Fig. 2d). Cells from the two datasets were mixed in most clusters, but overall the datasets occupied different parts of the two-dimensional space indicating subtle transcriptomic differences within cell types, potentially depicting differences in their developmental stages. With the exception of four clusters consisting primarily of cells from our neural samples or organoids, all other clusters contained cells originating from both systems. The clusters specific to the iCoMoNSC-derived cultures were composed of iCoMoNSCs (1 and 3), OPCs (14) and maturing neurons (18). The organoid-specific clusters 2 and 20 contained iPSCs, or neuroepithelial-like/retina progenitors (6), as well as cycling ventral and dorsal progenitors (11) (Fig. 2d,f). Overlapping neuronal populations included midbrain/hindbrain cells (cluster 7), intermediate progenitors and early cortical neurons/ventral progenitors (cluster 16), ventral progenitors and cortical neurons (clusters 0, 4, 8, 10 and 13). Overlapping glial subtypes were labeled as radial glia (clusters 5, 9, 12 and 15) and early radial glia/neuroepithelial-/neuroectodermal-like cells (cluster 17), whereas cluster 21 cells were annotated as choroid-plexus/mesenchymal-like cells. Altogether, our neural model intersected with human cortical brain organoids within clusters representing intermediate and ventral progenitors, early and late cortical neurons and radial glial cells. Differences were driven by the source cells, i.e., NSCs in our model and iPSCs in the brain organoid models. These results demonstrate that our neural model contained neuronal and glial cells of a transcriptional maturation level similar to that in brain organoids. Notwithstanding the differences in the source cells, these data indicate that, even in the absence of a 3D organ-like cell organization, neuronal and glial maturation in our iCoMoSCS-derived neural model is comparable to that in cortical organoids.

## Induced TDP-43 pathology in a minority of neurons

TDP-43 pathology characterizes affected brain regions of patients with TDP-43 proteinopathies, yet it was recently reported that only <2% of cortical cells show pathological changes in TDP-43 in FTLD^41^. To simulate this, we transduced young iCoMoNSC-derived neural networks with human TDP-43-HA-expressing lentiviral vector42 (Extended Data Fig. 7), with a titer designed to reach ~2% of transgenic cells in the network at 1 week of induction (Fig. 3a-b). Yet the percentage of transgenic neurons in the network gradually decreased (Fig. 3b), indicating that TDP-43-HA overexpression was toxic to human neurons. We then analyzed TDP-43-HA biochemically, using SarkoSpin, a method for the specific enrichment of pathological TDP-43 species, recently developed in our lab^18^. Interestingly, we detected progressive accumulation of TDP-43-HA in the SarkoSpin pellet, accompanied by fragmentation and appearance of high molecular weight bands and smear indicative of aggregation (Fig. 3c, Extended Data Fig. 8a), reminiscent of postmortem TDP-43 proteinopathy patient brains18. Specifically, in total cell lysates, we detected progressive fragmentation of full-length TDP-43-HA, to its 35 kDa and 25 kDa C-terminal fragments over time (Extended Data Fig. 8b). Both TDP-43 fragments accumulated in the SarkoSpin pellet, while full-length TDP-43-HA was present in the soluble fraction (SarkoSpin supernatant) at 2 weeks post induction, but was progressively redistributed to the SarkoSpin pellet in later time points (Fig. 3c). Collectively, our data demonstrate that overexpression of wild-type TDP-43-HA in human neurons resulted in progressive aggregation and fragmentation and the loss of TDP-43-HA-overexpressing cells. Surprisingly, while we found no evidence that TDP-43-HA overexpressing neurons develop TDP-43^p403/404^-positive inclusions (Fig. 3a), the latter emerged and amplified over time within non-transgenic neurons present in the same neural network (Extended Data Fig. 8c,d), showing that pathological TDP-43 changes extend beyond the initially affected cells. At early time points, TDP-43^p403/404^ signal appeared in the form of small, dot-like pre-inclusions that were largely confined within the soma (Extended Data Fig. 8e), whereas at later time points, these inclusions were larger and additionally present within neuronal processes (Extended Data Fig. 8f). This indicates progressive maturation of TDP-43 inclusions into aggregates in aged human neural networks.

**Figure 3.**
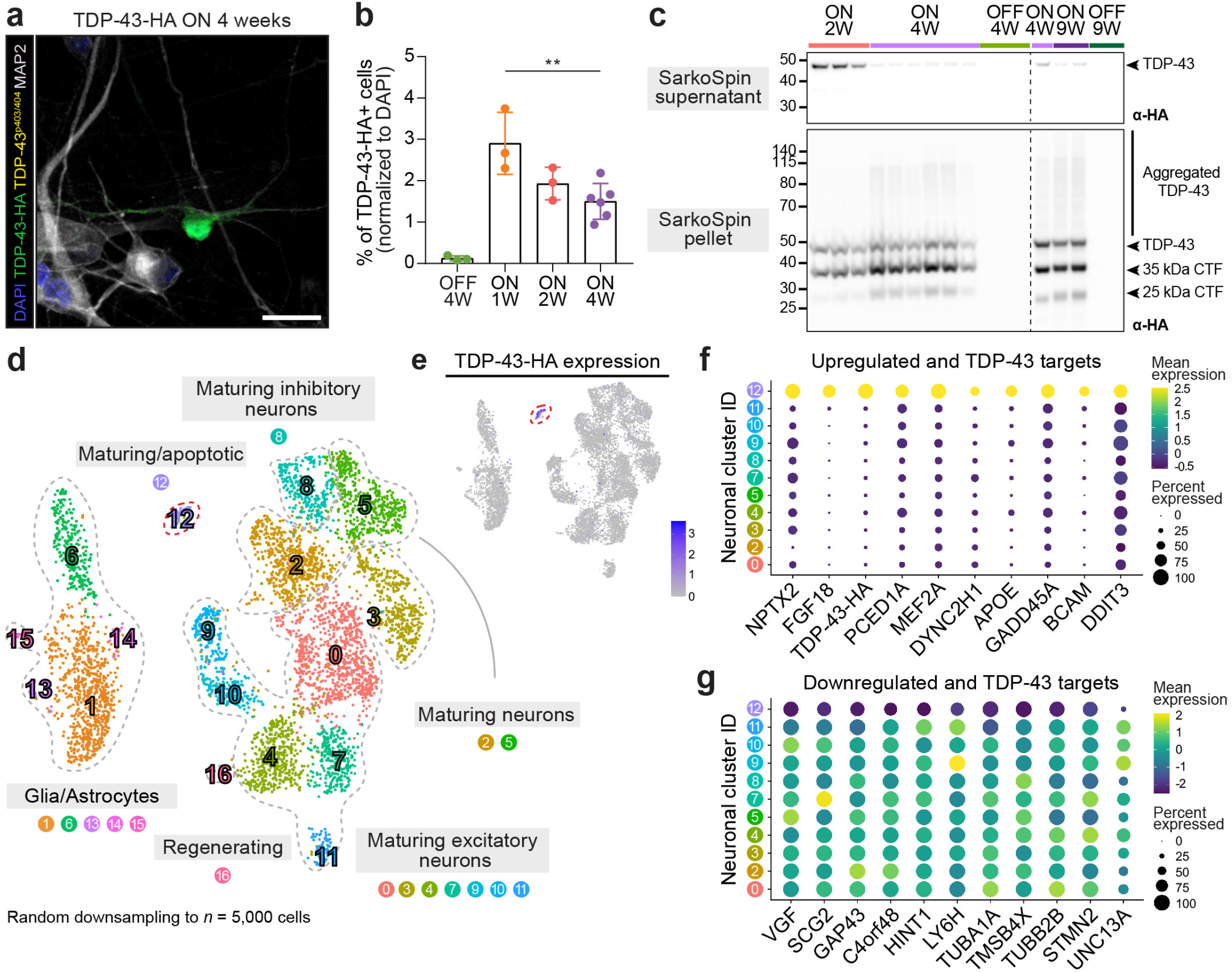
Distinct transcriptional profile in neurons with induced TDP-43 pathology. **a**, Immunofluorescence of TDP-43-HA in iCoMoNSC neurons. Scale bar, 25 μm. **b**, Quantification of TDP-43-HA-positive neurons over time. Each dot represents a biological replicate. **c**, Western blots of SarkoSpin supernatants (top) and pellets (bottom). Line separates independent experiments. **d**, UMAP of single-cell RNA-seq TDP-43 overexpression experiment. Colors indicate clusters, dashed lines highlight different cell types. Red dashed line depicts cluster 12, containing cells expressing TDP-43-HA. **e**, UMAP highlighting TDP-43-HA expression confined in cluster 12. **f**, Dot plot with the scaled average expression of the top cluster 12-upregulated and (**g**) −downregulated, including the significantly downregulated *UNC13A,* marker genes when compared to all other neuronal clusters.

## Distinct transcriptional profile in neurons with TDP-43 pathology

To understand the effect of sparse TDP-43-HA overexpression and related pathology in human neural networks, we induced its expression for 2 or 4 weeks, before harvesting the cells for scRNA-seq. Samples were analyzed at our middle stage (~3 months), consisting of both inhibitory and excitatory neurons (Fig. 2a,b) interconnected into functional networks (Fig. 1k). Between 6000-10000 cells per sample were retained after preprocessing and filtering with a median number of detected genes of 4300-5100 and median number of UMIs between 15000 and 20000. We identified 17 clusters (Fig. 3d) with a very similar cell type distribution (apoptotic, glial/astrocytic, maturing inhibitory and excitatory neurons) to our non-transduced middle-aged samples (Fig. 2a,b). We quantified the expression of the TDP-43-HA transcript in all samples using the scRNA-seq analysis pipeline Alevin^43^ and identified marker genes for each cluster. This analysis revealed a single cluster (number 12) that was almost exclusively composed of cells overexpressing TDP-43-HA (1.3% of cells in cluster 12 are from the OFF sample, 43.3% from ON 2W, 39.3% from ON 4W r1 and 16.1% from ON 4W r2) (Fig. 3e, Extended Data Fig. 8g). Cells in cluster 12 had an increase in total TDP-43 expression (log2FC 2.96), compared to all other neuronal clusters, and we were able to detect the transduced construct in 84.4% of cluster 12 cells, but only 2.6% of all other neuronal cells. Overexpression of TDP-43-HA over a period of 2 or 4 weeks altered the expression of several genes that were identified as cluster 12 markers and were either upregulated (Fig. 3f) or downregulated (Fig. 3g) when compared to all other neuronal clusters. Among the top 10 downregulated genes was *STMN2^3,4^* a previously known human-specific RNA target of TDP-43. *UNC13A5,6*, another recently discovered TDP-43 target was not among the top 10 markers, but was also significantly downregulated in TDP-43-overexpressing cells (Fig. 3g), validating the relevance of our model for human disease. Excitingly, several marker genes that have not yet been associated with or directly connected to ALS or FTLD neuropathology, but are linked to neuronal survival and/or death, have been found upregulated in cluster 12. Among those, Neuronal Pentraxin 2 (*NPTX2*), a neuron-specific gene, also known as neuronal activity-regulated protein (*NARP*), is secreted and involved in long-term neuronal plasticity. To understand if any of these RNAs were directly bound by TDP-43, we analyzed previously published iCLIP datasets from control and FTLD patient brain samples17. Interestingly, with the exception of *C4orf48* and *GADD45A*, all other cluster 12-misregulated RNAs are indeed TDP-43 binding targets and the binding is altered in FTLD compared to control patient brains (Fig. 4a).

**Figure 4.**
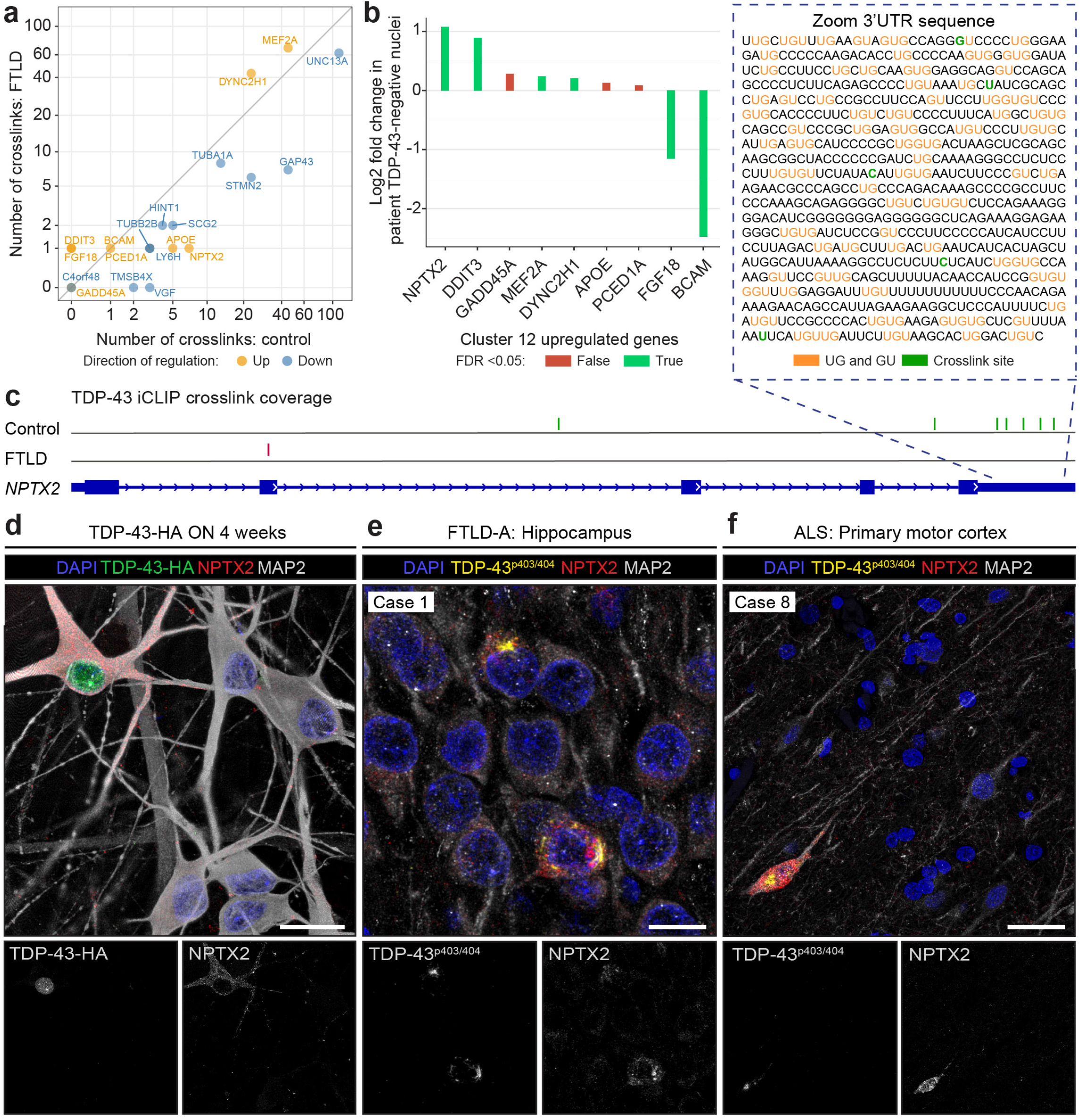
NPTX2 is misaccumulated in FTLD and ALS patients. **a**, Plot shows the number of iCLIP crosslinks^17^ in both FTLD patient and control brain samples in mRNA of the cluster 12 up- and - downregulated genes. **b**, Log2 fold change in RNA expression of the top 10 cluster 12-upregulated markers between TDP-43-negative and TDP-43-positive neuronal nuclei from FTLD patients ^44^. **c**, Location of iCLIP crosslink sites in *NPTX2* in both control (green) and FTLD patient (red) human brain^17^. Zoom-in: iCLIP crosslinks (green) and UG- or GU-repeats (orange). **d**, Only TDP-43-HA-overexpressing neurons showed detectable NPTX2 levels in immunofluorescence. **e**, Immunofluorescence of human brain sections: FTLD-A hippocampus and (**f**) ALS primary motor cortex demonstrating inclusion-like NPTX2 signal only in MAP2-positive neurons containing TDP-43^p403/404^-positive aggregates in the cytoplasm. Scale bars, (**d**) 20 μm, (**e**) 10 μm, (**f**) 30 μm.

## NPTX2 is misaccumulated in FTLD and ALS patients

We then investigated if any of our newly identified cluster 12 marker genes were altered in ALS and FTLD patient brain samples. To that end, we re-analysed the recently reported RNA-seq dataset comparing the transcriptomic profiles of single nuclei from individual FTLD-ALS neurons with (TDP-43-negative) or without (TDP-43-positive) nuclear clearance^44^, which is a consequence of pathological TDP-43 accumulation and sequestration of functional protein. Out of the nine cluster 12-upregulated RNAs, six were significantly altered, and four were significantly upregulated in TDP-43-negative nuclei. The strongest upregulation was in *NPTX2* mRNA, which was increased two-fold in TDP-43-negative FTLD-ALS patient neurons compared to controls (Fig. 4b). Moreover, iCLIP data^17^ analysis showed that TDP-43 directly bound *NPTX2* mRNA (Fig. 4a), primarily at its 3’UTR within a highly GU-rich region (Fig. 4c), the sequence specifically identified by the RNA recognition motifs of TDP-4313,45,46. Importantly, this TDP-43/*NPTX2* interaction was reduced in FTLD patient brains, as shown by the loss of iCLIP crosslinks, marking positions of direct protein-RNA interactions. Indeed, while in control brains, five TDP-43 crosslinks were identified on *NPTX2 3’UTR,* none were detected in FTLD brains (Fig. 4c). Collectively these data suggest that TDP-43 directly regulates *NPTX2* mRNA levels via binding on its 3’UTR, an event that is disturbed in human neurons with TDP-43 pathology.

To understand if NPTX2 protein is specifically altered in neurons with TDP-43 pathology, we then used immunofluorescence in our human neural networks and found that this normally secreted and synaptically-localized protein^47,48^ is aberrantly accumulated within neuronal somata and processes of TDP-43-HA-expressing neurons (Fig. 4d, Extended Data Fig. 9a). Validating our cellular model and analysis, affected neurons also depicted increase in MEF2A and decrease in STMN2 levels (Extended Data Fig. 9b-c), as we predicted (Fig. 3f-g). We then wondered if NPTX2 accumulation also occurred in patient brains (Supplementary Table 1). To this end, we used immunofluorescence for NPTX2, TDP-43^p403/404^ and MAP2 (neuronal marker) and found that while NPTX2 levels were very low in adult human neurons of patients, it accumulated into dense inclusions in both the somata and processes of neurons with TDP-43 pathology in the granule cell layer of the dentate gyrus of the hippocampus of both FTLD-TDP Type A and Type C^49,50^ patients as well as in the frontal cortex of FTLD-A patients and in the primary motor cortex of an ALS patient (Fig. 4e,f, Extended Data Fig. 10). Interestingly, NPTX2 did not appear to co-aggregate with TDP-43, suggesting that its loss of function, rather than a gain, is responsible for this novel NPTX2 pathology. Overall, our work identified aberrant NPTX2 accumulation as a direct consequence of TDP-43 misregulation in disease, suggesting a novel neurotoxic pathway for TDP-43 proteinopathies.

## Discussion

Long-term self-renewing NSC lines derived from ES or iPSCs provide an excellent source of expandable cells^7,8,51^ for creating models of neurogenesis and CNS-related diseases. We generated iCoMoNSCs and showed that they represent a uniquely homogenous population of self-renewing NSCs with the potential to give rise to mature and diverse neuronal and glial subtypes. Notably, only a very small percentage (~0.3%) of iCoMoNSCs was identified as committed neuroblasts. Upon terminal differentiation, iCoMoNSCs consistently generated mixed neuronal and glial cultures with electrophysiologically active neurons that formed interconnected networks with spontaneous coordinated activity. Importantly, scRNA-seq confirmed the progressive maturation of neuronal and glial subtypes comparable to that in human brain organoids^9^.

We then explored how overexpression of TDP-43 in a minority of neurons would affect these neural networks over time, aiming to simulate the situation in the diseased brain that reportedly contains only ~2% of cells with TDP-43 pathology^44^. After 4 weeks of TDP-43-HA overexpression, we observed a drop in soluble TDP-43-HA levels paralleled by progressive insoluble TDP-43 aggregation and C-terminal fragmentation^52^, as well as progressive loss of TDP-43-HA-positive cells. scRNA-seq revealed that cells with TDP-43 overexpression and pathology were characterized by a distinct transcriptional profile, with downregulation of *STMN2^3,4^* and *UNC13A^5,6^*, indicating loss of TDP-43 function. Importantly, this approach uncovered a novel set of up- and downregulated genes. Remarkably, with the exception of two, all were direct RNA targets of TDP-43 and their binding was previously found to be altered in the human FTLD brain^17^.

Contrasting our novel RNA targets with RNA-seq data from TDP-43-negative cortical neurons of ALS/FTLD patients^44^ highlighted *NPTX2* as the most significantly upregulated RNA in human patients. TDP-43 binds *NPTX2* mRNA at its 3’UTR, an event that is disturbed in FTLD^17^, suggesting a repressive role of this interaction. Importantly, the binding mode and increased *NPTX2* RNA levels indicate a regulation mechanism distinct from the previously described cryptic exon suppression^23^ recently shown for *STMN2^3^*^,4^ and *UNC13A^5^*^,6^. Notably, this misregulation led to aberrant accumulation of NPTX2 protein, which we consistently observed not only in our cellular model, but also in the brains of ALS and FTLD patients. Indeed, NPTX2 accumulation in patients reliably marked cells with TDP-43 pathology, without co-aggregation of the two proteins. Interestingly, NPTX2 was recently shown to be decreased in the CSF of symptomatic genetic FTLD patients, in a manner correlating with clinical severity^53^. Collectively, these data indicate that while global NPTX2 levels decrease with age^54^ and in dementia^53–56^, neurons with TDP-43 pathology have increased levels of NPTX2, likely due to loss of TDP-43 binding on *NPTX2* mRNA.

What could the consequences of this aberrant increase of NPTX2 levels in adult neurons be? NPTX2 is a neuronal pentraxin and an immediate early gene regulated by neuronal activity^48,57,58^. It is a Ca^2+^-dependent lectin^48^ that is secreted from presynaptic terminals^47,48^, where it regulates synaptogenesis and glutamate signaling via AMPA receptor (AMPAR) clustering^57,59,60^. NPTX2 binding to neuronal pentraxin receptor (NPTXR)^59^ mediates synaptic maintenance, plasticity and postsynaptic specialization^57,61^ in both excitatory and inhibitory synapses^61^. NPTX2 enhances glutamate receptor 1 function, and its upregulation causes synaptic remodeling through recruiting Ca^2+^-permeable AMPARs on the neuronal membrane^62^. Glutamate excitotoxicity has long been proposed to be one of the major pathological pathways selectively killing vulnerable neurons in ALS^63^ and other neurodegenerative diseases^64^. Since excitotoxicity is primarily propagated via Ca^2+^ influx through Ca^2+^-permeable receptors^65^, it is attractive to speculate that NPTX2 malfunction in neurons with TDP-43 pathology may increase their vulnerability to glutamate excitotoxicity in ALS and FTLD. We propose that NPTX2 may be a key neuronal target of TDP-43 and that its aberrant increase in affected neurons may trigger a pathological cascade leading to neuronal loss in ALS/FTLD. In this context, NPTX2 may represent an important therapeutic target for TDP-43 proteinopathies, along with the recently discovered STMN2 and UNC13A.

## Methods

### iPSC-derived, colony morphology neural stem cells (iCoMoNSCs)

Similarly to our previous study on human ESCs7, iPSC colonies were manually picked and partially dissociated into smaller clumps and transferred into non-adhesive culture dishes and induced to form embryoid bodies (EBs) in the presence of EB medium (iPS media without bFGF). After 5-7 days, EBs were transferred onto Poly-L-Ornithine (Sigma P4957-50ML; 20 μg/ml in sterile water Gibco 10977035, 1 hour at 37°C followed by 3 washes with PBS Gibco 10010015) / Laminin (Gibco 23017-015; 5 μg/ml in PBS, 1 hour at 37°C) coated dishes (P/L) and left to adhere in NSC media (DMEM/F12 Gibco 21331046; 0.5X B27-supplement Gibco 12587-010, 0.5X N2 supplement Gibco 17502-048; 1X GlutaMAX Gibco 35050-061; 25 ng/ml bFGF Gibco PHG0261). Formation of neural rosettes was observed within 4-10 days. Neural rosettes were manually dissected and picked under EVOS™ XL Core Imaging System (LifeTech AMEX1000), and after dissociation re-plated onto fresh P/L-coated dishes in NSC media. After 2-5 days, new and smaller rosettes appeared (R1 rosettes) with the presence of heterogeneous contaminating cells. The R1 rosettes were then manually dissected, picked and dissociated into smaller clumps and re-plated onto fresh P/L dishes. After further 2-5 days, new rosettes (R2) with minimal contaminating cells appeared. R2 rosettes were then routinely monitored to identify small groups of radially organized cells that were present outside of the neural rosettes and represented an independent “clone-like population”, the iCoMoNSCs. These small patches of iCoMoNSCs were then manually picked and transferred onto freshly P/L-coated 24 well plates. Clones that showed clear radial and consistent morphology, good attachment, survival and proliferation upon transfer were enzymatically detached using 0.05% Trypsin (Gibco 15400-054 diluted in PBS) and 1X Defined Trypsin Inhibitor (Gibco R-007-100) and expanded for numerous passages and banked in NSC freezing media (NSC media + 10% DMSO). Upon further expansion to P9-13, iCoMoNSCs lines were prepared for karyotype check at Cell Guidance Systems (according to CellGS fixed sample protocol). iCoMoNSCs clone 10/80 was used in the study. iCoMoNSCs are available upon request.

### Differentiation of iCoMoNSCs into human neural networks

iCoMoNSCs were plated in NSC media at 75 000 cells/cm2 onto Matrigel-coated (Corning #354234; ~0.15 mg/ml diluted in cold DMEM/F12 Gibco #11330032 and incubated at least 1 hour at 37°C) 6 well plates and left to recover and proliferate to reach ~95% confluency. At this point, NSC media was switched to “D3” differentiation media (DMEM/F12 Gibco #11330032; 0.5X B27+ supplement Gibco #17504-044, 1X N2 supplement Gibco #17502-048; 1X GlutaMAX Gibco #35050-061; 1X Penicillin/Streptomycin Sigma #P4333-100ML) supplemented with 5 μM Forskolin (Cayman #AG-CN2-0089-M050), 1 μM synthetic retinoid Ec23 (Amsbio #AMS.SRP002-2), 500 nM Smoothened agonist SAG (Millipore #5666600) for the first 5 days. On the days 6-10, Ec23 was increased to 2 μM. On days 11-25, Ec23 was decreased to 10ng/ml, SAG to 50nM and BDNF (PeproTech # 450-02), GDNF (PeptroTech # 450-10) and CNTF (Alomone labs #C-240) were added at 20 ng/ml. At day 26 and onwards, media was switched to maturation media (1:1 DMEM/F12:Neurobasal (Gibco #21103049) mix; 1X B27+ supplement, 1X N2 supplement; 1X GlutaMAX, 5 μM Forskolin, BDNF, GDNF, CNTF, NT-3 (PeproTech #450-03) and IGF-1 (Stem Cell #78022) all at 20 ng/ml and 10 μM cAMP. Media was changed daily and almost completely for the first 10 days, whereas from this point on only 2/3 of media was changed 3 times a week.

### Cloning of the AutoTetON cassette, lentiviral vector preparation and transduction

Using NEBuilder HiFi DNA Assembly Cloning Kit (NEB #E5520), human wild-type, full length TDP-43 with a C-terminal HA tag from a published pcDNA5 plasmid containing the human TDP-43 cDNA sequence^66^ was inserted into our autoregulatory, all-in-one TetON cassette (AutoTetON; build based on Markusic et. al 2005^42^), which was previously inserted into a pLVX lentiviral transfer vector (Clontech # 632164), while deleting CMV-PGK-Puro, generating Auto-TDP-43-HA lentiviral transfer vector. AutoTetON cassette was build using both NEBuilder and Q5 site directed mutagenesis (NEB #E0554S) kits and consists of the Tet-responsive promoter *P*_tight_, consisting of seven tet operator sequences followed by the minimal CMV promoter (sequence source pCW57.1; Addgene #41393), driving the inducible expression of the downstream TDP-43-HA, followed by downstream IRES2 sequence (sequence source Addgene #60857), which is immediately followed by T7 tag fused to SV40 NLS, which was fused to rtTA-Advanced (sequence source Addgene #41393), which made the rtTA predominantly nuclear, making it readily available for the system, while the T7-tag made rtTA a useful, independent marker of the transgenic cells. See Extended Data Fig. 7 for more details.

Auto-TDP-43-HA was then packaged into lentivirus (LV) via co-transfection with CMV-Gag-Pol (Harvard #dR8.91) and pVSV-G (Clontech, part of #631530) plasmids into production HEK293T cells adapted to grow in serum-free conditions (OHN media; based on Opti-MEM (ThermoFisher #11058-021), supplemented with 0.5% B27-(ThermoFisher #12587-010), 0.5% N2 (ThermoFisher #175020-01), 1% GlutaMAX (ThermoFisher #35050038) and bFGF (25ng/ml; ThermoFisher #PHG0261)), which reduces the expression of the GOI from the transfer vector (i.e. it eliminates traces of tetracyclines in the FBS) as well as eliminates serum-carry over into the LV supernatant. Medium was changed the following morning and supernatants were then collected 48 hours post transfection (36 hours post media change), centrifuged at (500g, 10 min, 4°C), filtered through Whatman 0.45μm CA filter (GE #10462100) and concentrated using Lenti-X™ Concentrator (Takara #631232) according to the producer instructions (overnight incubation). The resulting lentiviral pellets were then resuspended in complete neuronal maturation media to achieve 10x concentrated LV preparations, which were titrated using Lenti-X™ GoStix™ Plus (Takara #631280). 10x concentrate of Auto-TDP-43-HA LV was then used at 1600ng (of lentiviral p24 protein as per GoStix Value (GV)) per well of a 6 well plate of differentiated human neural cultures (~2 months old) along with 3μg/ml of polybrene (Sigma-Aldrich #TR-1003-G), pipetting the LV concentrate directly onto the culture (drop-wise). Complete neuronal maturation media was then added to reach 750μl total. Medium was exchanged completely the following day. TDP-43-HA expression was induced by 1μg/ml of Doxycycline (DOX; Clontech #631311) when needed.

### Patient post-mortem brain immunofluorescence

Formalin-fixed, paraffin-embedded hippocampal, frontal or primary motor cortex patient (ALS, FTLD-TDP Type A, FTLD-TDP Type C) sections (see **Supplementary table 1**) were used. All FTLD tissue samples were donated to Queen Square Brain Bank for Neurological Disorders at UCL Queen Square Institute of Neurology with full, informed consent. Anonymized autopsy ALS sample was collected by the Institute of Neuropathology at UZH. According to Swiss law, anonymized autopsy tissues do not fall within the scope of the Human Research Act and may be used in research. Sections were deparaffinized in three Xylene rounds (5 minutes each) and rehydrated in decreasing EtOH washes (2x 100% for 10 minutes; 2x in 95% for 5 minutes; 80, 70 and 50% for 5 minutes each) and finally submerged in MilliQ water for 10 minutes. Antigen retrieval was then performed by microwave heat treatment in sodium citrate buffer (0.01M, pH 6.0). Sections were then cooled down on ice for 10 minutes and once quickly washed in PBS, followed by 3x PBS washes for 5 minutes at RT before blocking with blocking buffer (BB; 5% normal donkey serum Sigma-Aldrich #S30-M, 3% BSA (Sigma #A4503) and 0.25% Triton-X100 (Sigma #T9284) in PBS for 30 minutes at RT. Primary antibodies (Supplementary Table 2) were then diluted in BB and 300 μl of the antibody mix was evenly put per slide/section and covered by parafilm. Slides were put into a wet and dark incubation chamber and incubated overnight at RT. Slides were rinsed once in PBS and then washed 3x 5 minutes in PBS on a shaker. Secondary antibodies (Supplementary Table 2) diluted in BB, centrifuged for 30 minutes at 15000 x g at RT and 500 μl of the antibody mix was evenly put per slide/section and put into the incubation chamber to incubate for 2.5 hours at RT. Slides were then rinsed once in PBS, washed 1x 5 minutes in PBS on a shaker and to stain the nuclei (DNA), 500 μl of DAPI solution (Thermo Scientific #62248; diluted to 1ug/ml in PBS) was evenly added onto the slide/section and incubated for 10 minutes at RT in the incubation chamber. Slides were rinsed once in PBS, washed 2x 5 minutes in PBS on a shaker, followed by Sudan Black (0.2% in 70% EtOH; Sigma #199664) autofluorescence quench for 10 minutes at RT on shaker. Slides were rinsed 6 times in PBS and left washing in PBS until mounted and coverslipped in ProLong Diamond Antifade Mountant (ThermoFisher #P36961) and then left to dry in the chemical cabinet overnight at RT in the dark. Mounted sections were stored at 4°C in the dark. Stained patient brain sections were imaged using Leica SP8 Falcon inverted confocal for high power, high resolution microscopy (63X oil objective; 1.7 or 3x zoom; 2096×2096 pixels at 0.059μm/pixel or 1848×1848 pixels at 0.033μm/pixel, respectively, approx. 20 z-steps per stack at 0.3μm). White laser and HyD detector settings were kept the same for each staining combination and all imaged conditions. Huygens professional (Scientific Volume Imaging, Hilversum, Netherlands) was used to deconvolute the stacks and the deconvoluted images were further post-processed in Fiji ^67^ to produce a 3D projection for data visualization (1st image of the 3D projection is shown in figures). Post-processing settings were kept the same for all images of all sections from all patients (except for DAPI and MAP2 channels for certain sections when higher brightness settings were used to reach intensity that allowed proper visualization).

### High-Density Microelectrode Arrays

CMOS-based HD-MEAs^35^ were used to record the extracellular action potentials of iCoMoNSC-derived human neural networks. The HD-MEA featured 26,400 electrodes, organized in a 120 × 220 grid within a total sensing area of 3.85 × 2.10 mm^2^. The electrode area was 9.3 × 5.45 μm^2^, and the center-to-center electrode distance (pitch) was 17.5 μm, which allows for recording of cell electrical activity at subcellular resolution. Up to 1,024 electrodes could be simultaneously recorded from in user-selected configurations. The HD-MEA featured noise values of 2.4 μVrms in the action-potential band of 0.3 - 10 kHz and hads a programmable gain of up to 78 dB. The sampling frequency was 20 kHz.

#### HD-MEA Recordings

The recording setup was placed inside a 5% CO2 cell-culture incubator at 37°C. Recordings were performed using the “Activity Scan Assay” and “Network Assay” modules, featured in the MaxLab Live software (MaxWell Biosystems AG, Zurich, Switzerland), as previously described^37^. The spontaneous neuronal activity across the whole HD-MEA was recorded using 6,600 electrodes at a pitch of 35 μm in 7 electrode configurations for 120 seconds. The most “active” 1,024 electrodes were then used to record network electrical activity for 300 seconds. Active electrodes were identified based on their firing rate, and, among those, the 1,024 electrodes featuring the highest firing rates were selected.

#### HD-MEA Metrics

We used metrics similar to those described in Ronchi et al., 2021^37^ to characterize and compare the neuronal cultures; we used network, single-cell and subcellular-resolution metrics. As network metrics (Extended Data Fig.6a) we used the burst duration (BD), inter-burst interval (IBI), inter-burst interval coefficient of variation (IBIcv)^37^. As single-cell metrics (Extended Data Fig.6b) we used the mean firing rate (MFR), mean spike amplitude (MSA), and the inter-spike interval coefficient of variation (ISIcv)^37^. Additionally, we included the following extracellular waveform metrics (Extended Data Fig.6c), extracted from SpikeInterface^38^, an open-source Python-based framework to enclose all the spike sorting steps:

1. Half width half maximum (HWHM), half width of trough of AP wave at half amplitude.
2. Peak-to-trough ratio (PTr), ratio of peak amplitude with respect to amplitude of trough.
3. Peak to valley (PtV), time interval between peak and valley.
4. Repolarization slope (RepS), slope between trough and return to baseline.
5. Recovery slope (RecS), slope after peak towards recovery to baseline.

As subcellular-resolution metrics, we extracted the action potential propagation velocity (Vel)^37]^ and the axon branch length (BL).

The percentage of active electrodes (AE) was also computed to measure the overall number of electrodes that could detect action potentials^37^.

#### HD-MEA Data Analysis

Data analysis was performed using custom-written codes in MATLAB R2021a and Python 3.6.10 and are available upon request.

Spike sorting was performed to identify single units in the extracellular recordings. We used the Kilosort2^68^ software within the SpikeInterface^38^ framework and the corresponding default parameters. We automatically curated the spike sorting output using the following parameters (<u>link</u>):

1. Inter-spike interval violation threshold (ISIt) = 0.5 The ISIt takes into account the refractory period, which follows every action potential (AP). The assumption is that if two APs occur within a too short time interval, they most likely come from two different neurons.
2. Firing rate threshold (FRt) = 0.05 The FRt sets the minimum firing rate of a neuron to be considered as “good” unit.
3. Signal-to-noise ratio threshold (SNRt) = 5 The SNRt takes into account the ratio between the maximum amplitude of the mean AP waveform and the noise characteristics of the specific channel.
4. Amplitude cutoff (ACt) = 0.1 The ACt takes into account the false-negative rate, thus the fraction of spikes per unit with an amplitude below the detection threshold.
5. Nearest neighbors hit rate (NNt) = 0.9 After computing the principal component for a unit, the NNt is used to check on the fraction of the nearest neighbors that fall into the same cluster.

#### HD-MEA Statistical Analysis

Statistical comparisons to compare samples from more than two populations were performed using the Kruskal-Wallis H test. In case the null hypothesis was rejected, we performed a post-hoc Dunn test with Sidák correction for multiple comparisons (Dunn-Sidák multiple-comparison test).

Statistical analysis was performed in MATLAB R2021a.

### scRNA sequencing of human neural networks

Duplicates (except for TDP-43-HA OFF and TDP-43-HA 2 weeks samples) of human neural networks (young (1.5 months), middle (3 months) and old (7.5 months)) and TDP-43-HA experiment samples at middle stage, were dissociated into single-cells suspension using Papain Dissociation System (Worthington #LK003150), passed through 70μm and 40μm cell strainers (Falcon #07-201-431 and #07-201-430), and resuspended in HIB++ media (Hibernate™-E Medium medium (Gibco #A1247601) supplemented with EDTA (1mM final; Invitrogen #AM9260G), HEPES (10mM final; Gibco #15630080), with 1X B27+ supplement (Gibco #17504-044), 1X N2 supplement (Gibco #17502-048); 1X GlutaMAX (Gibco #35050-061), BDNF (PeproTech #450-02), GDNF (Alomone labs #G-240), CNTF (Alomone labs #C-240), NT-3 (PeproTech #450-03) and IGF-1 (Stem Cell #78022) all at 20 ng/ml)) to 1000 cells per μl using CASY Cell Counter (Innovatis AG).

### Single-cell RNA sequencing (scRNA-seq) using 10X Genomics platform

The quality and concentration of the single-cell preparations were evaluated using an haemocytometer in a Leica DM IL LED microscope and adjusted to 1,000 cells/μl. 10,000 cells per sample were loaded in to the 10X Chromium controller and library preparation was performed according to the manufacturer’s indications (Chromium Next GEM Single Cell 3ʹ Reagent Kits v3.1 protocol). The resulting libraries were sequenced in an Illumina NovaSeq sequencer according to 10X Genomics recommendations (paired-end reads, R1=28, i7=8, R2=91) to a depth of around 50,000 reads per cell. The sequencing was performed at Functional Genomics Center Zurich (FGCZ).

### Single-cell data analysis

Data was processed with CellRanger for demultiplexing, read alignment to the human reference genome (GRCh38) and filtering to generate a feature-barcode matrix per sample. Cell doublets were removed with scDblFinder^69^ and outlier cells were detected and filtered with the “scater” R package^70^. In short, cells with more than 3 median-absolute-deviations away from the median number of UMIs, the number of features and the percentage of mitochondrial genes were removed. For the 7.5 months old samples, we additionally filtered cells with less than 2000 UMIs and less than 1500 detected features. For the TDP-43 overexpression experiment, we additionally filtered cells with less than 5000 UMIs and less than 2500 detected features.

Seurat v3^71^ was used for log-normalization and to identify the top 2000 highly variable genes per sample. Louvain clustering was always performed with resolution 0.4 based on a shared nearest neighbor graph constructed from the top 20 principal components (PC). The Uniform Manifold Approximation and Projection (UMAP)^72^ was always computed from the top 20 PCs. Cell cycle scores were computed with Seurat using a list of G2/M and S phase markers from Kowalczyk et al. 2015^73^. Marker genes that are upregulated in one cluster compared to any other cluster with a log-fold change of at least 2 were identified with the findMarkers function from the scran R package^74^, which runs pairwise *t*-tests and combines the results into a ranked list of markers for each cluster. Heatmaps with marker gene expression show scaled mean log-counts of all cells in each cluster.

The two iCoMoNSC samples were integrated with Seurat using canonical-correlation analysis (CCA)^75^, the data was scaled and number of UMIs and the percentage of mitochondrial UMIs was regressed out before clustering.

The scRNA-seq data from three different NSC lines^8^ was downloaded from ArrayExpress (accession number E-MTAB-8379) and integrated with our iCoMoNSCs using Seurat, the data was scaled and the number of UMIs was integrated out before clustering.

The human organoid (404b2 and H9) scRNA-seq data^9^ was downloaded from ArrayExpress (accession number E-MTAB-7552). Our iCoMoNSC and human neural cultures (1.5, 3 and 7.5 months old) were integrated with cells from human brain organoids using Seurat, the data was scaled and the number of UMIs and the percentage of mitochondrial UMIs were regressed out before clustering.

In the four samples of the TDP-43 overexpression experiment, we additionally quantified the expression of the TDP-43-HA transcript, the total TDP-43 transcripts (endogenous TDP-43 transcript and TDP-43 sequence of the lentiviral construct and the lentiviral construct (transcribed part without the TDP-43 sequence) with Alevin^43^ using the list of identified barcodes from CellRanger as whitelist. The counts were added to the filtered feature-barcode matrix from CellRanger. The total TDP-43 log2FC between cluster 12 and all other neuronal clusters was computed using the subset of cells with total TDP-43 UMI count > 0 (91.1% of cells in cluster 12 and 29.5% of all other neuronal cells).

Additional Methods are detailed in the Supplementary Information.

## Data availability

scRNA-seq data will be made publicly available upon publication.

## Code availability

All code for the scRNA-seq data analysis is deposited on github https://github.com/khembach/neural_scRNAseq as a workflowr project ^76^. All plots and results can be accessed as well: https://khembach.github.io/neural_scRNAseq.

## Acknowledgements

We gratefully acknowledge the support of the Swiss National Science Foundation Foundation (grants: 310030_192650, PP00P3_176966) and the National Centre for Competence in Research (NCCR) RNA & Disease and the Swiss Foundation for Research on Muscle Diseases to MPo. M.H.P. was supported by the Milton Safenowitz Postdoctoral Fellowship from the ALS Association (16-PDF-247), postdoctoral fellowship from the University of Zurich (FK-15-097) and the Promotor-Stiftung from the Georges and Antoine Claraz Foundation. A.H. was supported by the European Research Council (ERC) Advanced Grant 694829 ‘neuroXscales’ and the corresponding proof-of-concept Grant 875609 ‘HD-Neu-Screen’. V.I.W. is supported by the FEBS Long-Term Fellowship and D.B. by the Czech Science Foundation (GACR grant no. 18-25429Y). T.L. is supported by an Alzheimer’s Research UK Senior Fellowship and Queen Square Brain Bank for Neurological studies is supported by the Reta Lila Weston Institute for Neurological Studies. The funders had no role in the experimental design, data collection, analysis, and preparation of the manuscript.

The authors would like to thank Emilio Yángüez, Andreia Joao Cabral de Gouvea and Doris Popovis from FGCZ for discussion and full support with single-cell RNA sequencing experiments and Ge Tan and Daymé González Rodríguez from FGCZ for bioinformatic support. We also thank Gery Barmettler from Center for Microscopy and Image Analysis (ZMB) UZH for technical help with TEM. All light imaging was performed with equipment maintained by the ZMB UZH.

## Author contributions

Conceptualization of the study was carried by M.H.P., K.M.B., S.R. and M.Po. M.H.P. generated iPSCs, iCoMoNSCs and differentiated them into neural networks, carried out cloning and lentiviral vector preparation, biochemistry and cell and human brain immunofluorescence as well as imaging and data analysis and prepared samples for calcium imaging, patch clamp, MEA and single-cell RNA sequencing experiments. K.M.B. analyzed the single-cell RNA sequencing data and re-analysed iCLIP and single nuclei RNA sequencing datasets. S.R. cultured neural cultures on the MEAs, performed MEA recordings, developed MEA metrics and analysed all recorded MEA data. V.I.W. performed human brain immunofluorescence and imaging and prepared samples for single-cell RNA sequencing. Z.M., E.M.H., and M.Pa. performed cell immunofluorescence and imaging, F.L. contributed to biochemistry experiments and S.S. performed synaptoneurosome isolation and western blots. V.H., and I.D. performed and analysed whole cell patch-clamp recordings. A.V.D.B. performed and analysed two photon calcium imaging. K.F., A.A. and T.L. provided human brain sections, neuropathological consultation and critical input on the study. D.B. provided critical feedback on the iCoMoNSC generation and characterization. M.D.R. provided feedback on sequencing data analyses. M.D.R., T.K., M.M., A.H., M.H.P. and M.Po. provided supervision. M.H.P., K.M.B., S.R., V.I.W. and M.Po. wrote and edited the manuscript and prepared figures. M.Po. directed the entire study. All authors read, edited, and approved the final manuscript.

## Additional information

Supplementary Information is available for this paper.

Correspondence and requests for materials should be addressed to

## Extended Data Figures legends

**Extended Data Figure 1.**
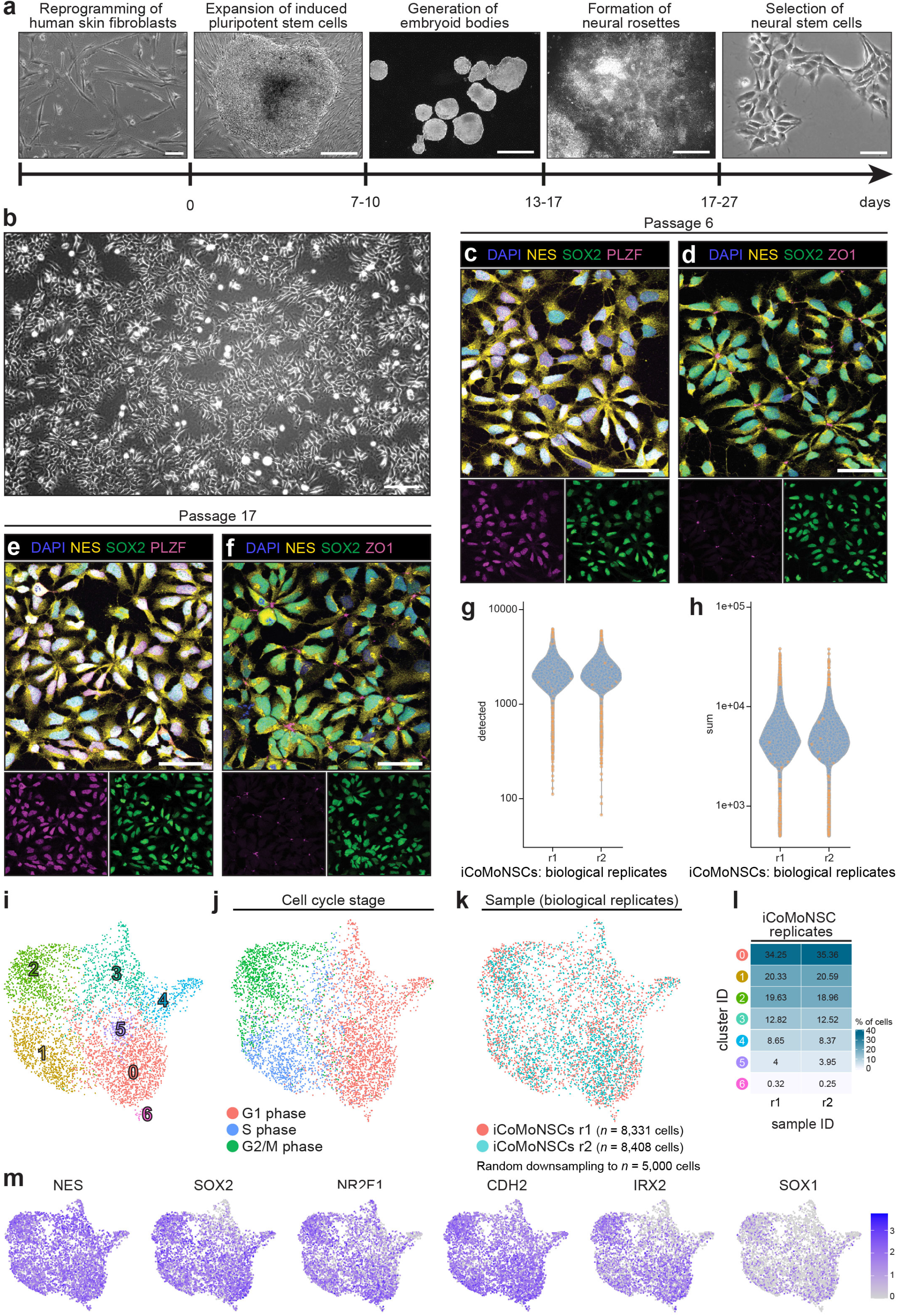
iCoMoNSCs characterization. **a**, Phase contrast images from different stages of iCoMoNSC generation. Human fibroblasts (left) were reprogrammed into iPSC colonies (2nd left), which formed embryoid bodies (middle) and then generated neural rosettes (2nd right). Patches of morphologically distinct colonies migrated out of rosettes and were isolated as clones (right). **b,**Phase contrast image showing the overall homogeneous morphology and pinwheel growth organisation of the iCoMoNSCs. NSC-marker immunofluorescence of iCoMoNSCs at early passage 6 (**c, d**) and late passage 17 (**e, f**) with NES, SOX2 and PLZF (**c, d**) or ZOI (**e, f**). Violin plots showing the number of genes (**g**) or UMIs (**h**) detected in all iCoMoNSCs by scRNA-seq in two biological replicates. **i**, UMAP of two integrated replicate samples of iCoMoNSCs clustering into 7 clusters, (**j**) same UMAP representing cell cycle stages or (**k**) showing the biological replicates whose distribution across the clusters is shown in (**l**). **m**, UMAP with normalized expression of selected genes across all iCoMoNSCs. Scale bars, (**a**) from left to right: 100 μm, 500 μm, 500 μm, 500 μm, 50 μm, (**b**) 150 μm, (**c-f**) 50 μm.

**Extended Data Figure 2.**
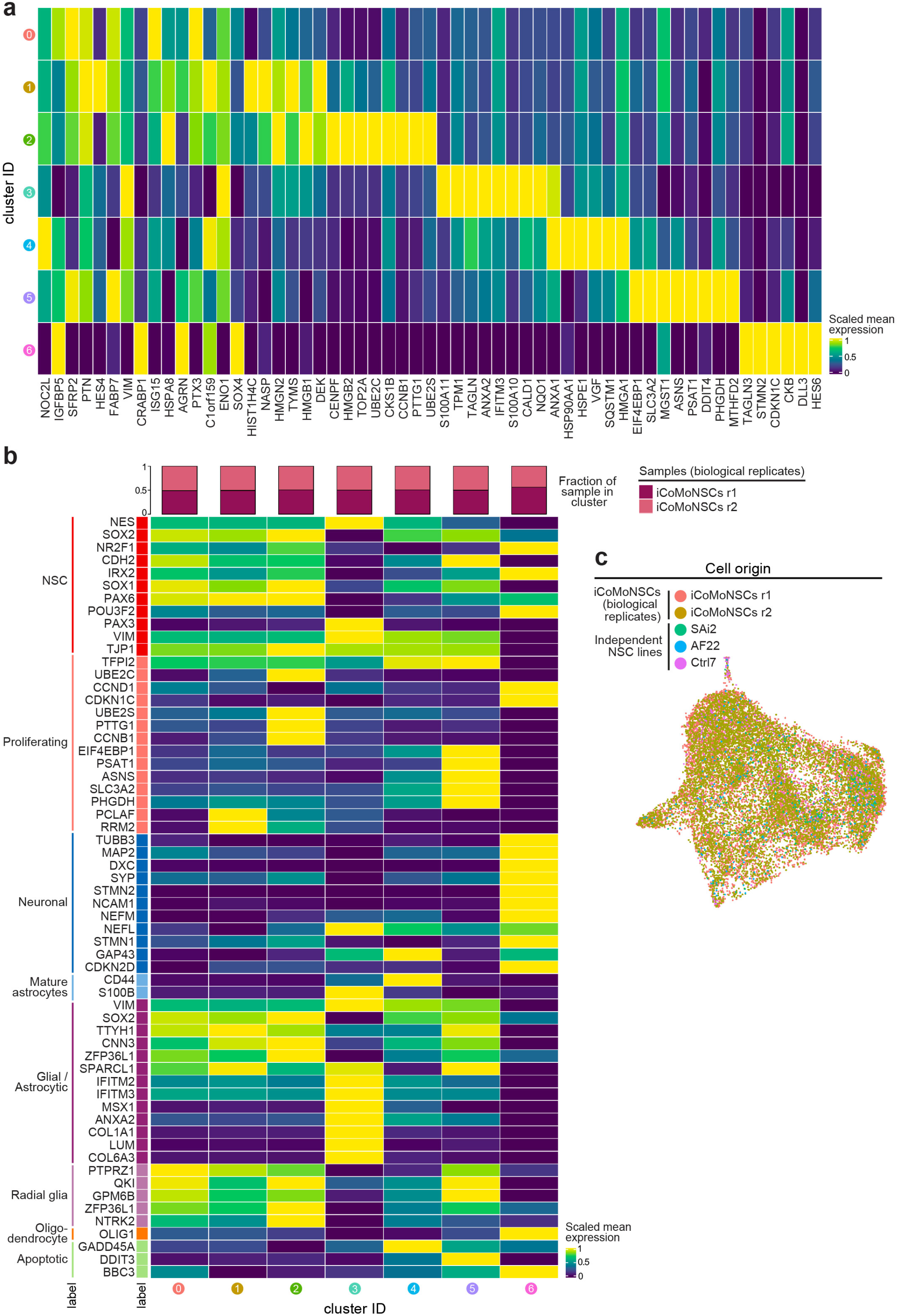
Cluster markers and known marker genes of iCoMoNSCs. **a**, Heatmap showing top cluster markers of iCoMoNSCs clusters. **b,**Heatmap showing gene expression of a set of known markers amongst the iCoMoNSCs clusters. **c**, UMAP of iCoMoNSCs integrated with 768 cells from three human neural stem cell lines (NES) showing individual samples in different colors.

**Extended Data Figure 3.**
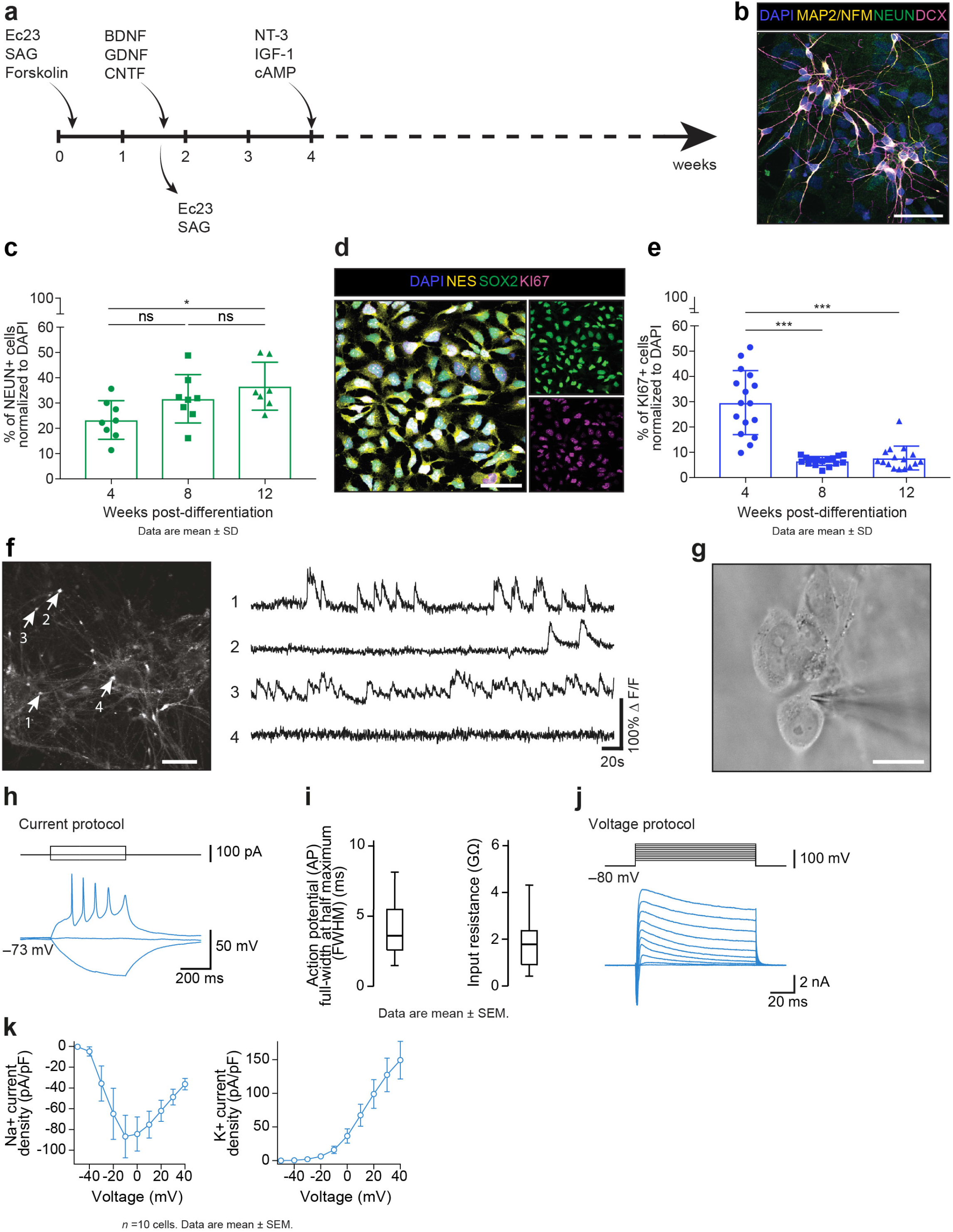
Differentiation of iCoMoNSCs. **a,**Schematics of iCoMoNSCs differentiation protocol. **b**, Immunofluorescence of neuronal markers MAP2/NFM, NEUN and DCX at 4 weeks of differentiation. **c**, Quantification of NEUN-positive cells over time in differentiation reaching approx. 40% at 12 weeks. Each data point represents an independent field of view normalized to DAPI^+^ nuclei. **d**, KI67 immunofluorescence of iCoMoNSCs at passage 6. **e**, Quantification of KI67-positive cells over time in differentiation showing a drop to 5% at 12 weeks. Each data point represents an independent field of view normalized to DAPI^+^ nuclei. **f**, 2-photon calcium imaging of neural networks bolus loaded with the AM ester form of Oregon Green BAPTA-1 showed typical firing neurons (neurons 1. and 3.). **g**, Infrared DIC image of whole-cell patch clamped human neuron at 4 months. **h**, Example voltage responses of a neuron elicited by tonic current injection. **i**, Average action potential and input resistance. **j**, Example current evoked by voltage steps from –60 mV to +40 mV (duration, 100 ms) from a holding voltage of –80 mV. Data recorded from the neuron in (**g, h**). **k,**Peak Na+ and K+ current densities plotted vs. voltage. Scale bars, (**b**,**d**) 50 μm, (**f**) 100 μm, (**g**) 20 μm.

**Extended Data Figure 4.**
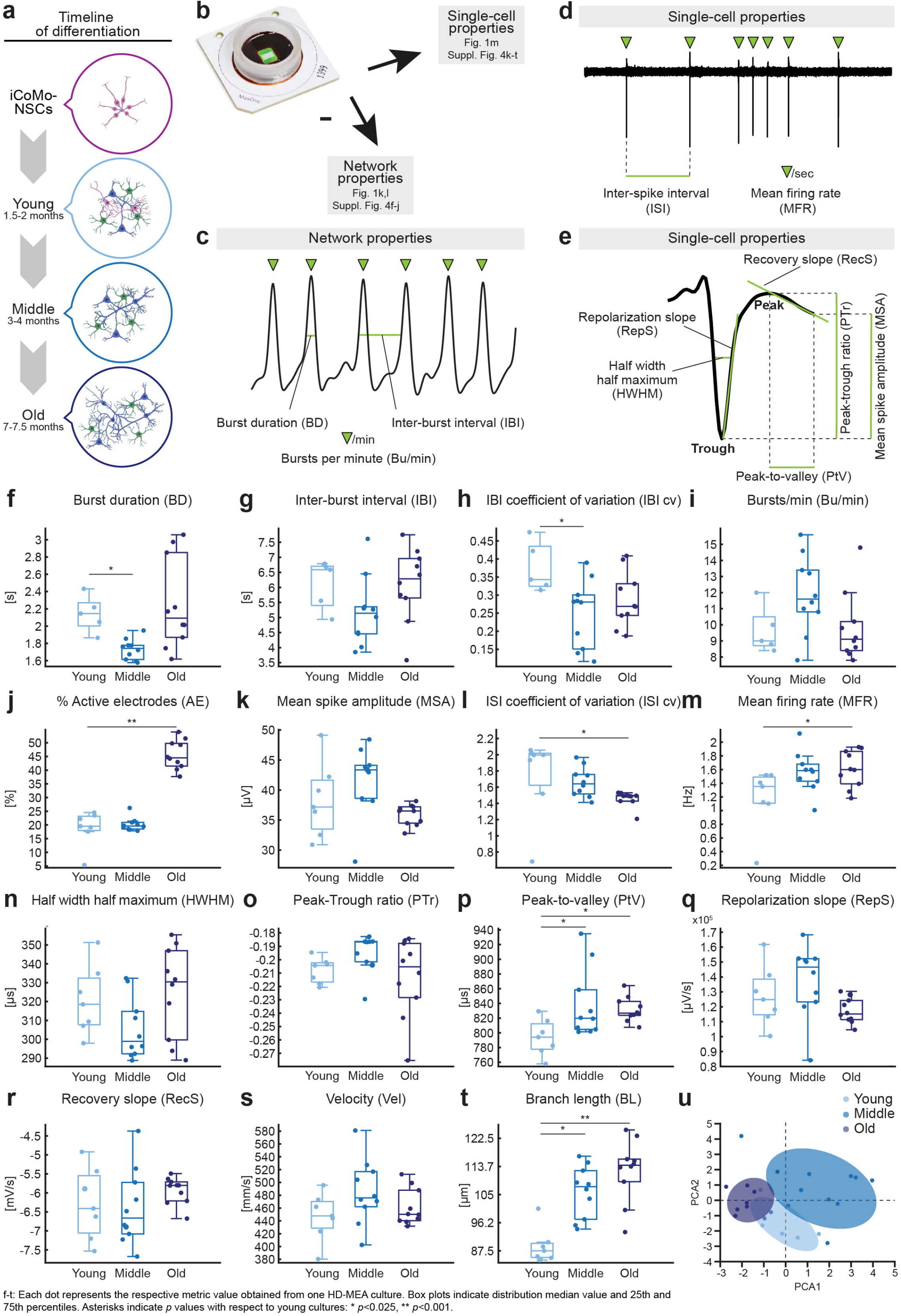
Schematics of network and single-cell properties and individual plots from data shown in figure 1l-n. **a**, Timeline of the experiment: starting from iCoMoNSCs (0 months), neural cultures were differentiated, sequenced and plated on MEAs at young (1.5 months), middle (3 months) and old (7 months) stages. **b**, HD-MEA chip mounted on a printed circuit board featuring a cell culture chamber (ring) and the microelectrode array in the center (green). Scale bar, 4 mm. **c**, Representative spike time histogram recorded from 1,020 electrodes used to illustrate how network metrics were extracted. **d**, Representative spontaneous APs recorded from one electrode, used to illustrate how single-cell metrics were extracted. **e**, Representative action potential (AP) used to illustrate how features were extracted from the AP shape. **f-t**, Box plots representing all results illustrated in **Fig. 1l, m**.

**Extended Data Figure 5.**
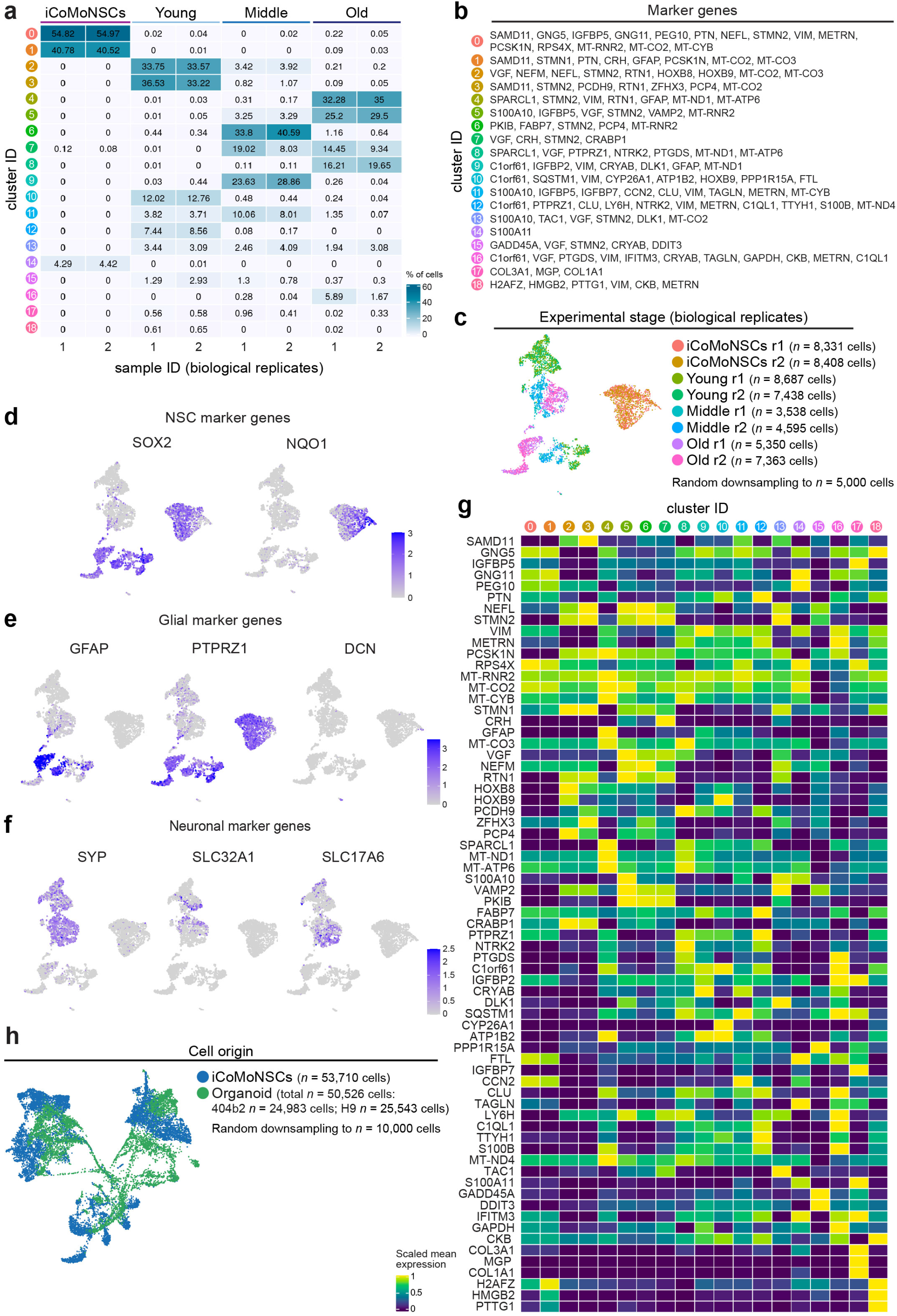
Neuronal and glial maturation of iCoMoNSC-differentiated human neural cultures. **a**, Heatmap of cell distribution from all experimental stages (in biological duplicates) amongst all clusters. **b**, Top cluster marker genes for each cluster. **c**, UMAP of iCoMoNSCs and young, middle and old human neural cultures with two replicates each showing cells from individual samples in different colors. **d**, UMAP with normalized expression of selected NSC, (**e**) glial and (**f**) neuronal marker genes across all samples. **g**, Heatmap with the expression of the top cluster markers from (**b**) from our aging experiment. **h,**UMAP of young, middle and old iCoMoNSC-derived neural cultures integrated with organoid datasets highlighting the cell origin.

**Extended Data Figure 6.**
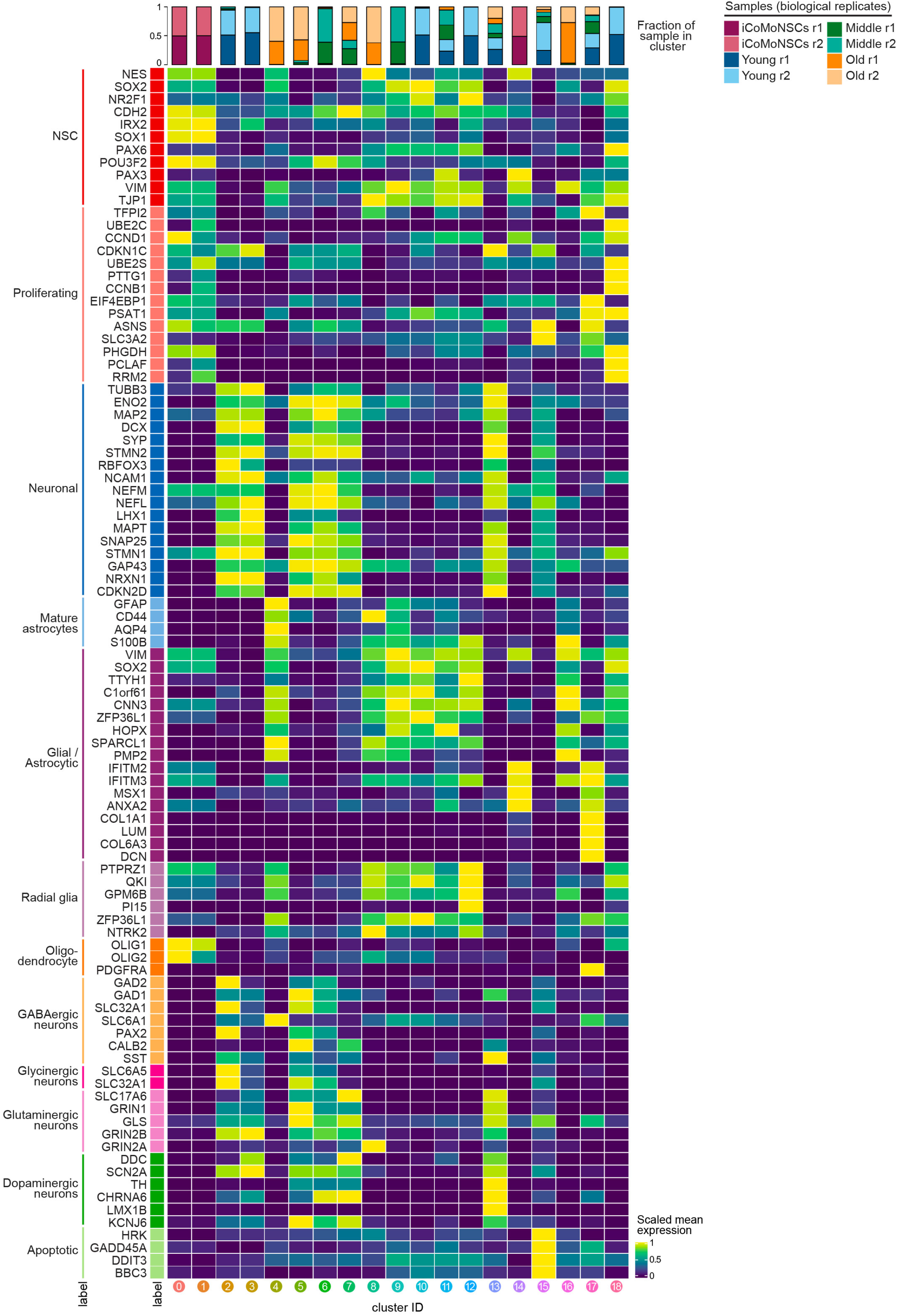
Heatmap with the gene expression of known marker genes amongst all clusters from our aging experiment.

**Extended Data Figure 7.**
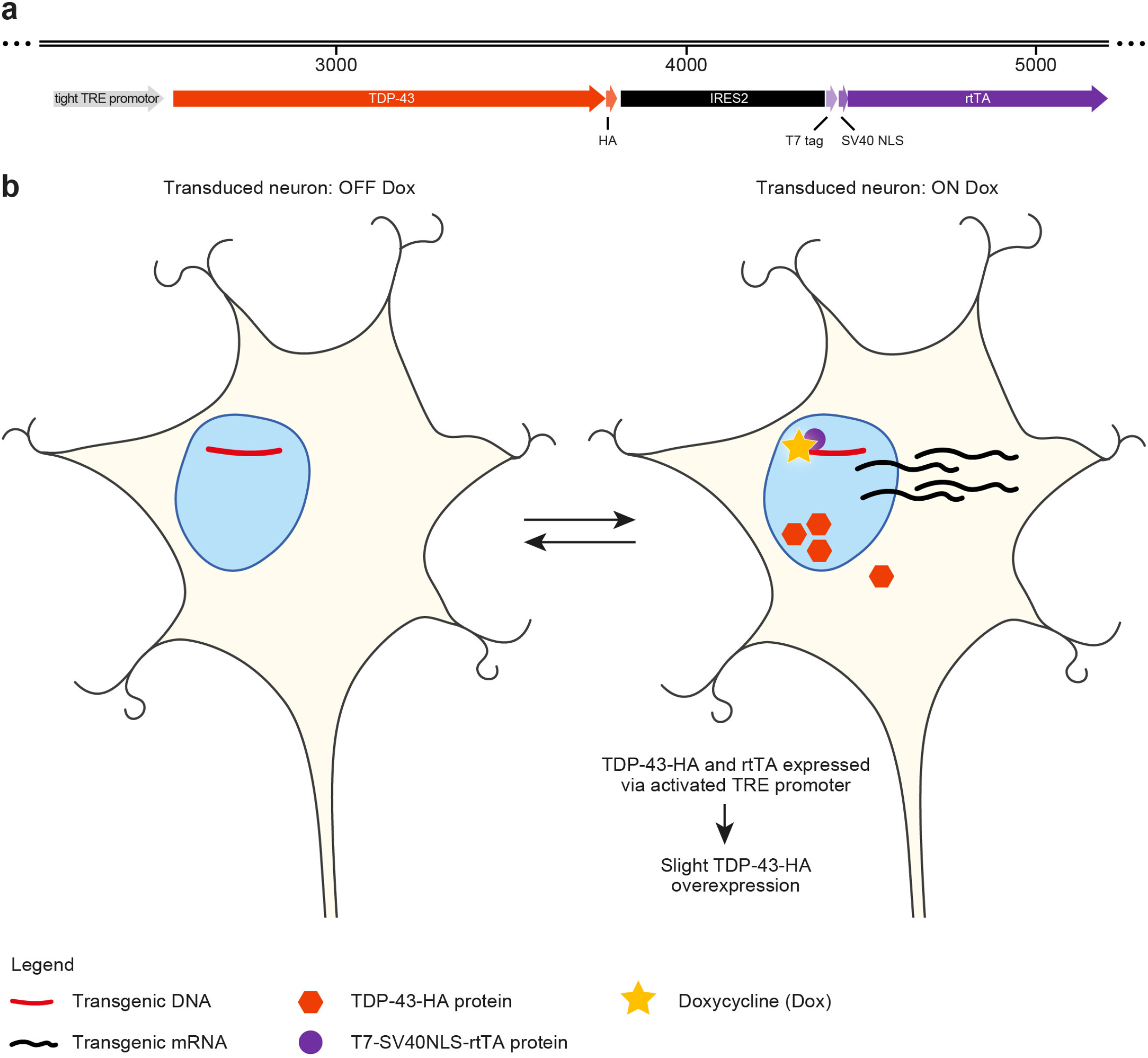
Autoregulatory, all-in-one TetON cassette and its function. **a**, Schematics of our improved, autoregulatory, all-in-one TetON cassette, which consists of a single tight TRE (*P*tight) promoter driving inducible expression of TDP-43-HA, which is linked via IRES2 to T7-tagged rtTA with SV40 NLS fused to its N-terminus to increase its nuclear localisation, making it a nuclear marker of transgenic cells. **b**, Schematics showing DOX OFF stage (left) in which no transgenic mRNA or protein could be detected. Upon addition of doxycycline (DOX ON stage; right), the nuclear, T7-tagged rtTA binds the Tet operators in the TRE promoter, which induces overexpression of TDP-43-HA as well as it leads to steady expression of rtTA needed for the whole cassette to function. The system is reversible.

**Extended Data Figure 8.**
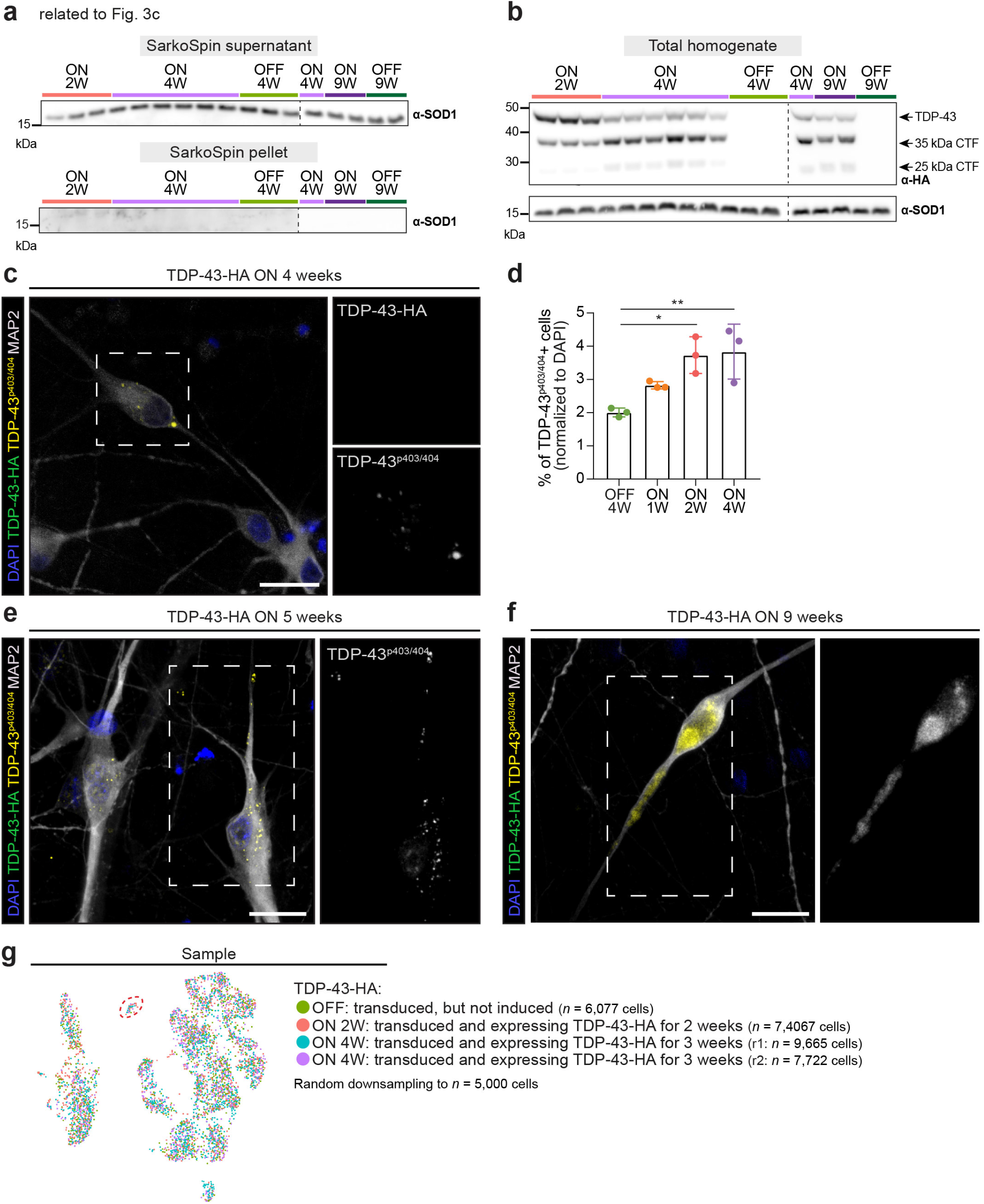
TDP-43-HA-induced pathology in human neural networks. **a,**SOD1 Western blot (WB) of SarkoSpin fractions of human neural networks overexpressing TDP-43-HA for 2, 4 or 9 weeks. **b**, WB on total homogenates (SarkoSpin input). SOD1 was used as a loading control. **c**, Immunofluorescence with phospho-specific (S403/404) anti-TDP-43 antibody revealed inclusion-like structures in the soma of TDP-43-HA-negative neurons. **d,**Quantification of TDP-43p^403/404^-positive cells over 4 weeks of TDP-43-HA overexpression. TDP-43^p403/404^-positive inclusions were later found localized to neurites at 5 weeks (**e**) and occasionally grew into-aggregate like structures at 9 weeks of TDP-43-HA overexpression (**f**). **g,**UMAP highlighting the four different samples of the single-cell RNA-seq TDP-43-HA experiment. Scale bars, 25 μm.

**Extended Data Figure 9.**
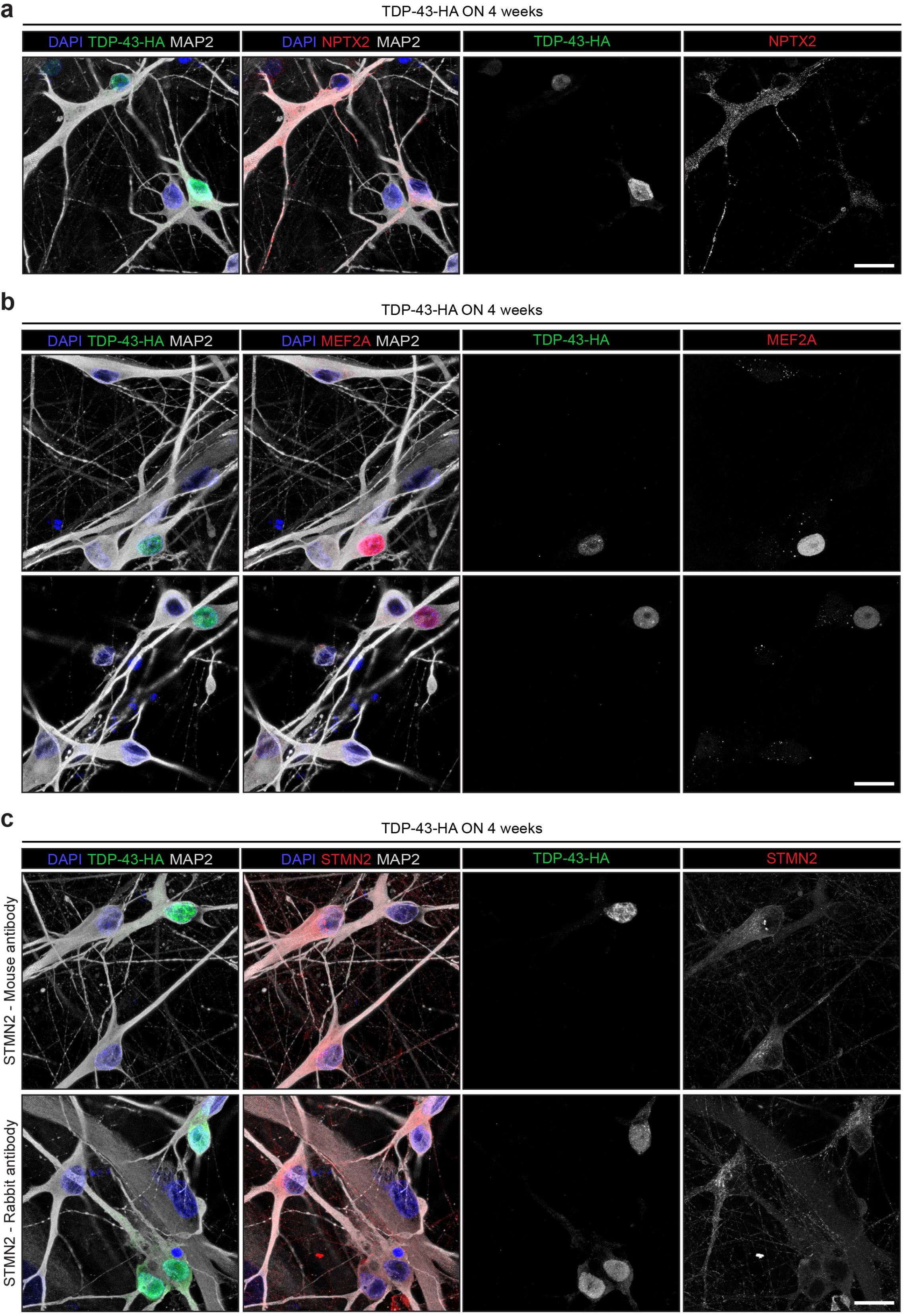
Immunofluorescence of cluster 12 misregulated genes in human neurons overexpressing TDP-43-HA. **a,**Immunofluorescence of human neural networks transduced to overexpress TDP-43-HA for 4 weeks demonstrating that only the TDP-43-HA-positive neurons expressed elevated levels of (**a**) NPTX2 in the soma and neurites, (**b**) MEF2A in the nucleus and (**c**) lowered levels of STMN2 in the soma and processes as demonstrated with 2 different antibodies. Scale bars, 20 μm.

**Extended Data Figure 10.**
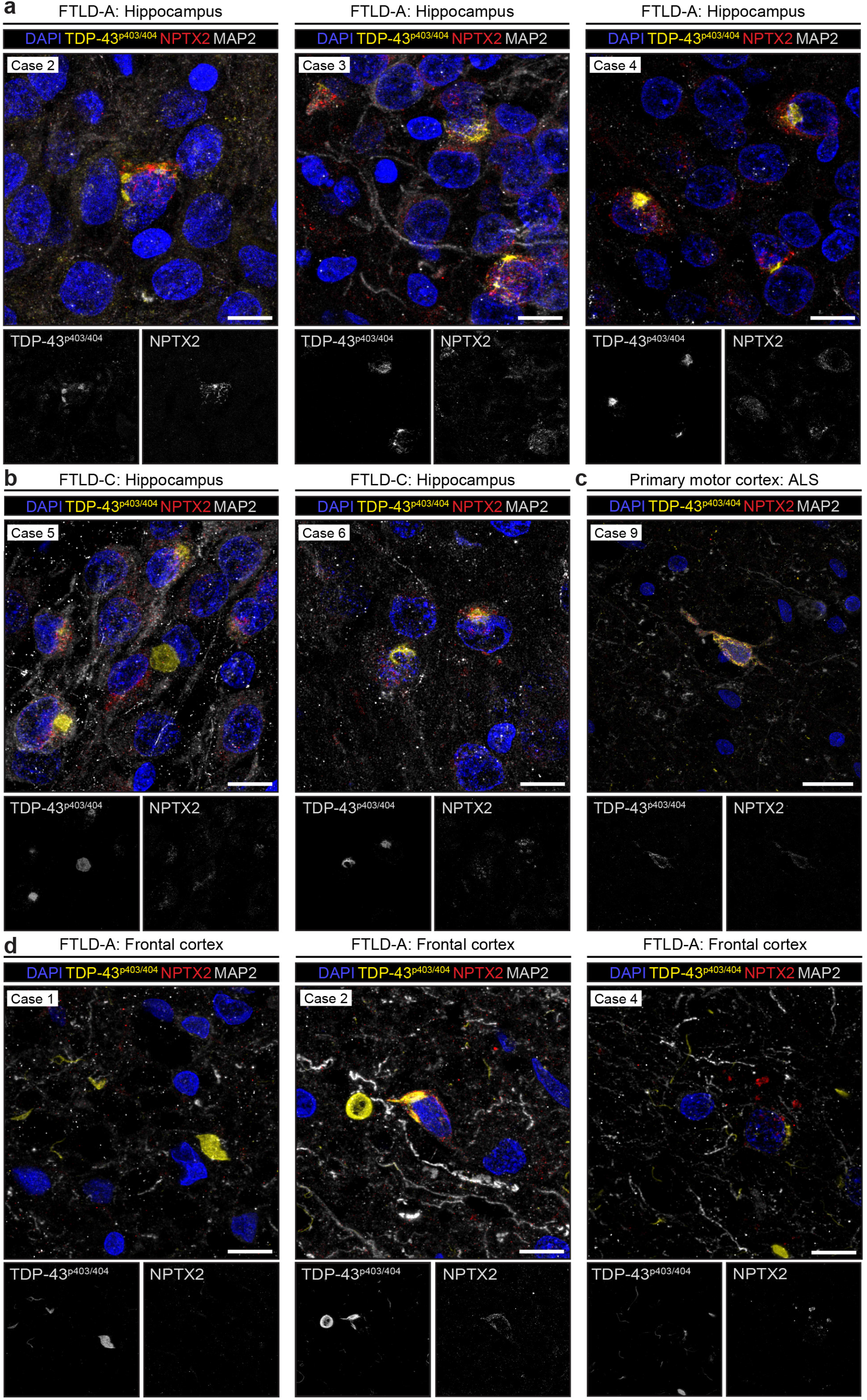
NPTX2 immunofluorescence in FTLD-A, FTLD-C and ALS patient brain sections. Additional immunofluorescence of multiple FTLD and ALS cases demonstrating NPTX2 accumulation and inclusions in neurons with TDP-43^p403/404^-positive aggregates in the hippocampus in (**a**) different FTLD-A cases, (**b**) different FTLD-C cases and in the primary motor cortex of an ALS case (**c**). **d**, Note that while the NPTX2 pathology could be observed in the cortex of both FTLD and ALS patients, it is only present when the TDP-43^p403/404^-positive aggregate-containing neurons could still be identified by the MAP2 staining (middle and right) as opposed to the disintegrated, “ghost neurons” that left behind only the TDP-43^p403/404^-positive aggregates (left). Scale bars, (**a**,**b**,**d**) 10 μm, (**c**) 20 μm.

