## Supplementary Information for "Human neural networks with sparse TDP-43 pathology reveal NPTX2 misregulation in ALS/FTLD"

**Supplementary methods**

**Generation of induced pluripotent stem cells (iPSCs)**

iPSCs were generated from control human early neonatal dermal fibroblasts (Gibco # C0045C) via episomal reprogramming using plasmids coding for Oct3/4, Sox2, Klf4 and p53-shRNA[1,2]. Plasmids (Addgene #27077, #27078 and #27080; 1 µg each) were electroporated into 1 million fibroblasts using 100 µl Neon tips (Invitrogen # MPK10025) on the Neon Transfection System (Pulse Voltage: 1650 V, Width: 10 ms, Number: 3; Invitrogen # MPK5000). After electroporation, cells were grown on gelatin-coated plates (EmbryoMax® 0.1% Gelatin Solution Millipore #ES-006-B) in fibroblast medium (DMEM Sigma #D5671; 20% FBS Gibco #10270-106; 1X GlutaMAX Gibco #35050-061; 1X NEAA Gibco #11140-050; 1X Penicillin/Streptomycin Sigma #P4333-100ML). On day 3 post electroporation (and then until day 12), 0.5 µg/ml of sodium butyrate (Sigma #B5887) was added to the culture media. The fibroblasts were then sub-cultured onto mouse embryonic fibroblasts (MEFs) (Global Stem # GSC-6001G or Millipore #PMEF-CFX) on the day 5 in iPSC medium (KnockOut™ DMEM/F-12 Gibco #12660012; 20% KnockOut™ Serum Replacement Gibco # 10828028, 1X GlutaMAX Gibco #35050-061; 1X NEAA Gibco #11140-050; 80 ng/ml bFGF Gibco #PHG0261; 1X EmbryoMax® 2-Mercaptoethanol Millipore #ES-007-E). The first iPSC colonies were observed around day 11. Positive live alkaline phosphatase staining (Invitrogen #A14353) was performed on day 14. iPSC colonies were individually manually picked on day 14-21, re-suspended and re-plated onto fresh MEF 24-well plates. iPSC clones were then expanded and the ones that displayed best proliferation and morphology (NiPS 8, 9 and 10) were banked at passage 10/11 in iPSC freezing media (iPSC media + 10% DMSO Sigma #D2650-100ML). Upon further expansion to P12-15, cells were prepared for karyotype check at Cell Guidance Systems (according to CellGS fixed sample protocol) and based on the karyotyping results, NiPS 10 was selected for further experiments.

**Preparation of human synaptoneurosomes (SNS)**

Neural cultures were washed with PBS and homogenized as they were scrapped off with the SNS buffer containing 0.35 M sucrose pH 7.4, 10 mM 4-(2 hydroxyethyl)-1-piperazineethanesulfonic acid (HEPES; Biosolve #08042359), 1 mM ethylenediaminetetraacetic acid (EDTA; VWR #0105), 0.25 mM dithiothreitol (Thermo Fisher Scientific #R0861), 30 U/ml RNAse inhibitor (Life Technologies #N8080119) and cOmplete-mini EDTA free protease inhibitor cocktail (Roche #11836170001; at least 2500 µl volume of buffer per one full 6 well plate of human neural cultures). Homogenates were passed sequentially through three 100 µm Nylon net filters (Millipore #NY1H02500), followed by one 5 µm filter (Millipore #SMWP013000). The final filtrate was re-suspended in 3 volumes of SNS buffer without sucrose and centrifuged at 2000 x g, for 15 mins at 4°C to yield a pellet containing synaptoneurosomes. The pellet was resuspended in 100 µl of SNS buffer, which was used for electron microscopy (60 µl) and western blot analysis (40 µl).

**Transmission electron microscopy (TEM)**

SNS pellet was prepared as mentioned above and submitted to the imaging facility (ZMB) of UZH. Briefly, SNS pellets were resuspended in 2X fixative (5% Glutaraldehyde in 0.2 M Cacodylate buffer) and fixed at RT for 30 mins. Samples were then washed twice with 0.1 M Cacodylate buffer before embedding into 2% Agar Nobile. Post-fixation was performed with 1% Osmium 1 h on ice, washed three times with ddH2O, dehydrated with 70% ethanol for 20 mins, followed by 80% ethanol for 20 mins, 100% for 30 mins, and finally Propylene for 30 mins. Propylene: Epon Araldite at 1:1 was added overnight followed by addition of Epon Araldite for 1 hour at RT. Sample was then embedded via 28 h incubation at 60 °C. The resulting block was then cut into 60 nm ultrathin sections using an ultramicrotome. Ribbons of sections were then put onto the TEM grid and imaged on the TEM - FEI CM100 electron microscope.

**Fixation, immunofluorescence and imaging of human neural cultures**

Neural cultures were fixed with pre-warmed 16% methanol-free formaldehyde (Pierce #28908) pipetted directly into the culture media, diluting it to 4% final, and incubated for 15 min at room temperature. Cells were then washed once with PBS (Gibco # 10010015) for 10 min, once with PBS with 0.2% Triton-X-100 (TX; Sigma #T9284) washing buffer (WB) for 10 min and then blocked with 10% normal donkey serum (Sigma-Aldrich #S30-M) and 0.2% TX in PBS blocking buffer (BB) filtered via stericup (Millipore #S2GPU02RE) for 30 min at room temperature (RT). Primary antibodies (Supplementary Table 2) were then diluted in BB and left incubated overnight (ON) at 4°C on an orbital shaker. Cells were then washed 3 x 15 min in WB at RT and secondaries (Supplementary Table 2) were then diluted in BB and incubated for 1.5 hours at RT. Cells were then again washed 3 x 15 min in WB at RT with DAPI (Thermo Scientific #62248) diluted to 1ug/ml in the final WB wash. Cells were finally washed 1 x 15 min in PBS at RT, and PBS were then added to the wells to store the stained cells at 4°C.

Stained human neural cultures were imaged using GE InCell Analyzer 2500 HS widefield microscope for quantification (40X air objective; 2D acquisition; 182 fields of view per well; 50µm separation to avoid counting cells twice) or with Leica SP8 Falcon inverted confocal for high-power, high-resolution microscopy (63X oil objective; 2096x2096 pixels at 0.059µm/pixel, approx. 20-30 z-steps per stack at 0.3µm). Laser and detector settings were kept the same for each staining combination and all imaged conditions. Huygens professional (Scientific Volume Imaging, Hilversum, Netherlands) was then used to deconvolute the stacks and the deconvoluted images were further post-processed in Fiji[3] to produce a flattened 2D pictures (Z-projection) for data visualization. IF experiments were repeated twice performing an independent experiment with biologically independent samples.

For quantification of NeuN- and KI67-positive cells, images were acquired using Leica SP5 inverted confocal with 63X oil objective (1024x1024 pixels at 1.7x zoom) and cells were counted and normalized to the total number of cells (total number of DAPI-positive nuclei).

Widefield image quantification was done using trained ilastik[4] algorithms to segment the pixels (positive vs background) of TDP-43-HA, TDP-43$^{p403/404}$ and DAPI stainings. Segmented pictures (182 images per well, per channel) were then exported and the total number of DAPI-positive nuclei quantified in Fiji[3] via batch processing using a custom macro, which is available upon request. TDP-43-HA or TDP-

43$^{p403/404}$-positive neurons were counted manually. Statistical analysis was performed in Prism (GraphPad San Diego, CA, USA) and one-way ANOVA followed by Tukey's multiple comparison test was applied on the datasets.

**Biochemical analysis of protein aggregates - SarkoSpin**

Modified SarkoSpin protocol[5,6] was used. Neural cultures were collected in 200 µl of cell lysis buffer (0.5% sarkosyl (Sigma #L5125-100G) + 0.5 µl Benzonase (Millipore #E1014-5KU) and 2 mM $MgCl_2$ in 1X HSI buffer (20 ml of 1X HSI: 10 mM TRIS, 150 mM NaCl, 0.5 mM EDTA, 1 mM DTT + 2 tablets of cOmplete EDTA free protease inhibitor cocktail (Roche 11873580001) and 1 tablet of PhosSTOP (Roche 0490684500). Re-suspended samples were transferred into low protein binding Eppendorf tubes. From the obtained total cell lysate, 30 µl was used for total blots, 5 µl was used for protein quantification and the remaining sample was further processed and analyzed by SarkoSpin. To 170 µl of the remaining cell homogenate, 178 µl 2X HSI with 4% sarkosyl and 52 µl 1X HSI (to obtain final sarkosyl concentration of 2% in total 400µl volume) was added. The samples were incubated at 37°C, 600 rpm for 45 min (Thermomixer, Eppendorf), and after increasing the volume by addition of 200 µl 1X HSI, the samples were vortex mixed and centrifuged at 21,200 x g on a benchtop centrifuge (Eppendorf) for 30 minutes at room temperature. Supernatants (~450 µl) were transferred to a new tube and the remaining supernatant was carefully removed to leave the pellets completely dry. The latter were then re-suspended in 80 µl of 0.5% sarkosyl 1X HSI and analyzed by immunoblot as described below. SarkoSpin was independently performed twice using biologically independent samples.

**SDS-PAGE and western blotting**

For total cell lysates and lysates of SNS pellets (of human neural cultures), protein concentration was adjusted using Pierce™ 660nm Protein Assay Reagent (#22660, all reagents from ThermoFisher unless otherwise stated) and 10 and 20 µg (total lysates and SNS pellets, respectively) of total protein per lane was used for immunoblots. Samples were re-suspended in final 1X LDS loading buffer (NP0007) with 1X final Bolt sample reducing agent (B0009), denatured at 70ºC for 10 mins, and loaded on Bolt 12% (SNS) or 4-12% (total and SarkoSpin fractions) Bis-Tris gels (NW04122BOX, NW04127BOX, NW00122BOX, NW00125BOX) using Bolt MES running buffer (B0002) and Bolt Antioxidant (BT0005) and run at 115V. Gels were transferred onto nitrocellulose membranes using iBlot 2 Transfer NC Stacks (IB23001) with iBlot 2 Dry Blotting System (IB21001) using P0 transfer method. Membranes were then blocked with 5% w/v non-fat skimmed powder milk in 0.05% v/v Tween-20 (Sigma P1379) in PBS (milk PBST) and probed with primary antibodies (Supplementary Table 2) overnight in PBST with 1% w/v milk, washed three times with PBST, followed by incubation with HRP-conjugated secondaries (Supplementary Table 2) in 1% milk PBST. After three washes in PBST, immunoreactivity was visualized by chemiluminescence using SuperSignal West Pico or Femto Chemiluminescent Substrate (PierceNet 34077, 34096) on Amersham Imager 600RGB (GE Healthcare Life Sciences 29083467). SarkoSpin western blots were independently performed twice using biologically independent samples.

**Two-photon calcium imaging**

Human neural cultures were bolus loaded with the AM ester form of Oregon Green BAPTA-1[7] by application (5 µM solution in HBSS with $Ca^{2+}/Mg^{2+}$ prepared from 500 µM stock solution[8] according to manufacturers' instructions. Shortly, human neural cultures that had been in differentiation for 12 weeks were briefly washed twice with pre-heated HBSS and then the 5µM of Oregon Green BAPTA-1 solution was added and incubated for 1 hour at 37°C. Cells were then briefly washed twice with pre-heated HBSS and subsequently allowed to stabilize at RT for 30 mins before imaging. Two-photon calcium imaging was performed on a Scientifica Hyperscope with a Ti:sapphire laser (Coherent; ~120 fs laser pulses) tuned to 840nm. Fluorescence images of 256x256 pixels at 4.25 Hz were collected with a 16x water-immersion objective lens (Nikon, NA 0.8). Data acquisition was performed by ScanImage.

Calcium imaging data were imported and analyzed using custom-written routines in ImageJ and MATLAB, which are available upon request. First, fluorescence image time-series for a given region were concatenated. The concatenated imaging data was then aligned using TurboReg to correct for small x-y drift (alignment on OGB-1 image series, NIH ImageJ; based on Thévenaz et al. 1998[9]). As a next step, the background was subtracted as the bottom first percentile fluorescence signal of the entire image. Average intensity projections of the imaging data were used as reference images to manually annotate regions of interest (ROIs) corresponding to individual neurons. Calcium signals were expressed as the mean pixel value of the relative fluorescence change $\Delta F/F = (F–F0)/F0$ in a given ROI. F0 was calculated as the bottom 10th percentile of the fluorescence trace.

**Electrophysiology**

Patch-clamp recordings were made on a SliceScope 2000 microscope (Scientifica Ltd, UK) using a HEKA EPC10/2 USB amplifier (HEKA Elektronik, Reutlingen, Germany). Data were low-pass filtered at 10 kHz and sampled at 200 kHz. Patch pipettes were pulled from borosilicate glass (Science Products, Hofheim, Germany) using a P-97 Puller (Sutter Instruments, Novato, CA). Pipettes had open-tip resistances of 4–8 MΩ when filled with intracellular solution containing (in mM): K-Gluconate 150, NaCl 10, HEPES 10, Mg-ATP 3, Na-GTP 0.3, BAPTA 0.2. A liquid junction potential of +13 mV was corrected for. The extracellular solution contained (in mM): NaCl 135, HEPES 10, Glucose 10, KCl 5, $CaCl_2$ 2, $MgCl_2$ 1. Series resistance ranged between 7–22 MΩ and was compensated online by 50–80% with 10 µs lag. All recordings were performed at room temperature (23–25°C). Resting membrane potential was measured in current-clamp after establishing the whole-cell configuration. To examine action potential firing and membrane properties, current injection protocols with incrementing amplitude (duration, 500 ms; step, ±10 pA) were applied from a holding voltage of approx. −80 mV (achieved by tonic current injection). Input resistance was calculated from the steady-state voltage deflection elicited by hyperpolarizing current injections of –10 pA. Membrane time constant was estimated from exponential fits to the voltage decay.

To record voltage-dependent ionic currents, cells were voltage-clamped at a holding potential of –80 mV. Currents were elicited by 100-ms voltage steps from –80 mV to +40 mV in steps of 10 mV and corrected for leak and capacitance currents using the P/5 method. For analysis of currents, cells were included if remaining series resistance was <10 MΩ. Data were analyzed with custom-written routines in Igor Pro (Wavemetrics, Lake Oswego, OR), which are available upon request.

**HD-MEA plating**

HD-MEAs were sterilized for 40 minutes in 70% ethanol. HD-MEAs were washed three times with sterile deionized water ($dH_2O$) and then dried. After sterilization, HD-MEAs were soaked in 20 µL of poly-D-lysine solution (P6407, Sigma-Aldrich) (diluted to 50 mg/mL in $dH_2O$) for 1 hour at room temperature to render the surface more hydrophilic; each HD-MEA was then washed three times with sterile $dH_2O$ and dried. Thereafter, HD-MEAs were coated with 10 µL Matrigel (354234, Corning Inc.), previously diluted in a plating medium (see below) at a 1:10 ratio and incubated at 37°C for 2 hours. Cells were centrifuged at 188 G-Force for 5 min, supernatant was aspirated, and plating medium was added to the cell pellet to reach the desired cell density (200'000 cells/HD-MEA). Matrigel was aspirated from HD-MEAs and cells were plated in a 10 µL drop on the HD-MEAs. 0.9 mL of medium was added after 2 hours of incubation at 37°C. 50% of plating medium was exchanged one day post plating and subsequently twice a week. Plating and culture medium was almost identical to maturation media (see methods) except that it used BrainPhys (05790, Stem Cell Technologies, Vancouver, Canada) as a base medium and was further supplemented with 1% Penicillin/Streptomycin (P4333-100ML, Sigma-Aldrich) to provide protection to cells that had to be periodically moved from culture incubator to recording culture incubator.

**Re-analysis of iCLIP data**

FASTQ files of human brain iCLIP[10] (three controls and three FTLD patients) were downloaded from ArrayExpress (E-MTAB-530)[11]. The data was re-analysed using the GRCh38 reference genome similarly than in the original publication with the iCount pipeline[12] for preprocessing, mapping and crosslink site identification. We merged the crosslink sites in the three control replicates and in the three FTLD replicates, summing the number of cDNAs per crosslink position. The two resulting crosslink files (control and FTLD) were used to prepare figure 4a and c in R version 4.0.5. All code has been deposited on github: https://github.com/khembach/Tollervey-iCLIP .

**Re-analysis of RNA-seq data from sorted nuclei**

We downloaded the RNA-seq gene count table as a supplementary data file (GSE126542_NeuronalNuclei_RNAseq_counts.txt.gz) from GEO (accession GSE126542). A differential gene expression analysis between TDP-43 negative and positive nuclei was performed in R version 4.0.5 with edgeR[13] using the quasi-likelihood framework[14] (empirical Bayes quasi-likelihood F-test). Reported false discovery rate (FDR) values are adjusted for multiple comparisons using the Benjamini-Hochberg method. The code is included in the scRNA-seq github repository https://github.com/khembach/neural_scRNAseq.

### Supplementary Methods References

**Supplementary Table 1 Demographics of cases used in the present study**

| Cases | Neuropathological diagnosis | Mutation | Gender | Age at disease onset | Age at death | Disease duration (years) | Post-mortem delay (h:min) |
|---|---|---|---|---|---|---|---|
| 1 | FTLD-TDP, Type A | | F | 57 | 63 | 6 | 85:20 |
| 2 | FTLD-TDP, Type A | C9orf72 | M | 51 | 61 | 10 | 35:15 |
| 3 | FTLD-TDP, Type A | C9orf72 | F | 56 | 67 | 11 | 85:35 |
| 4 | FTLD-TDP, Type A | | M | 59 | 70 | 11 | 44:05 |
| 5 | FTLD-TDP, Type C | | M | 64 | 74 | 10 | 19:00 |
| 6 | FTLD-TDP, Type C | | M | 71 | 76 | 5 | 39:30 |
| 7 | FTLD-TDP, Type C | | F | 58 | 73 | 15 | 37:55 |
| 8 | ALS | | F | 62 | 62 | 0.58 | 46:00 |

*ALS* amyotrophic lateral sclerosis, *C9orf72* Chromosome 9 open reading frame 72, *F* female, *FTLD-TDP* frontotemporal lobar degeneration with TDP-43 proteinopathy, *h:min* hours:minutes, *M* male

**Supplementary Table 2 Antibody list**

**Primary antibodies**

| Name | Species, Source | WB dilution | Cell IF dilution | Brain IF dilution |
|---|---|---|---|---|
| AQP4 | Rb, Novus Biologicals #NBP1-87679 | - | 1:200 | - |
| DCX | Gt, Santa Cruz Biotechnology #sc-8066 DCX | - | 1:2000 | - |
| GFAP | Gt, Abcam #ab53554 | - | 1:500 | - |
| HA | Rb, Cell Signaling Technology #3724 | 1:2500 | 1:500 | - |
| HA | Ms, Biolegend #901516 | - | 1:1000 | - |
| HA | Ms, ThermoFisher #26183 | - | 1:500 | - |
| KI67 | Rb, Abcam #ab16667 | - | 1:250 | - |
| MAP2 | Ms, Sigma #M1406 | - | 1:250 | - |
| MAP2 | Ch, Abcam #ab5392 | - | 1:1000 | 1:1000 |
| MEF2A | Rb, Santa Cruz Biotechnology #sc-17785 | - | 1:1000 | - |
| NEFL | Ms, Thermo Scientific #13-0700 | - | 1:2000 | - |
| Nestin | Ch, Online antibodies #ABIN187958 | - | 1:100 | - |
| NEUN | Ch, Millipore #ABN91 | - | 1:1000 | - |
| NPTX2 | Rb, Proteintech #10889-1-AP | - | 1:200 | 1:100 |
| NUMA | Rb, Bethyl #A301-510A | - | 1:200 | - |
| PLZF | Rb, Santa Cruz Biotechnology #sc22839 | - | 1:200 | - |
| PSD-95 | Ms, Abcam #ab2723-100 | 1:2000 | - | - |
| SNAP-25 | Ms, #SMI81 | 1:1000 | 1:500 | - |
| SOD1 | Rb, Enzo #ADI-SOD-100 | 1:15000 | - | - |
| SOX2 | Gt, Santa Cruz Biotechnology #sc17320 | - | 1:250 | - |
| STMN2 | Ms, Proteintech #67204-1-Ig | - | 1:100 | - |
| STMN2 | Rb, Proteintech #10586-1-AP | - | 1:200 | - |
| SYP | Rb, Santa Cruz Biotechnology #sc-9116 | 1:500 | 1:200 | - |

| | | | | |
|---|---|---|---|---|
| TDP-43$^{p403/404}$ | Ms, custom-made | - | 1:500 | 1:500 |
| VIM | Ch, Millipore #AB5733 | - | 1:2000 | - |
| ZO1 | Rb, Millipore #AB2272 | - | 1:500 | - |
| β-ACTIN | Ms, Sigma #A5441 | 1:5000 | - | - |

**Secondary antibodies**

| Name | Source | WB dilution | Cell IF dilution | Brain IF dilution |
|---|---|---|---|---|
| Donkey anti-Ch 488 | Jackson Immuno Research #JAC703-546-155 | - | 1:500 | - |
| Donkey anti-Ch 568 | Jackson Immuno Research #JAC703-586-155 | - | 1:500 | - |
| Donkey anti-Ch 647 | Jackson Immuno Research #JAC703-606-155 | - | 1:500 | - |
| Donkey anti-Gt 488 | ThermoFisher #A11055 | - | 1:500 | - |
| Donkey anti-Gt 594 | ThermoFisher #A11058 | - | 1:500 | - |
| Donkey anti-Gt 647 | ThermoFisher #A21447 | - | 1:500 | - |
| Donkey anti-Ms 488 | ThermoFisher #A21202 | - | 1:500 | - |
| Donkey anti-Ms 568 | ThermoFisher #A10037 | - | 1:500 | 1:400 |
| Donkey anti-Ms 647 | ThermoFisher #A31571 | - | 1:500 | - |
| Donkey anti-Rb 488 | ThermoFisher #A21206 | - | 1:500 | 1:400 |
| Donkey anti-Rb 568 | ThermoFisher #A10042 | - | 1:500 | - |
| Donkey anti-Rb 647 | ThermoFisher #A31573 | - | 1:500 | - |
| Goat anti-Ch 647 | ThermoFisher #A21449 | - | 1:500 | 1:400 |
| Goat anti-Ms-HRP | Jackson Immuno Research #115-035-146 | 1:5000 | - | - |
| Goat anti-Rb-HRP | Jackson Immuno Research #115-035-144 | 1:10000 | - | - |

*Ch* chicken, *Gt* goat, *HRP* horseradish peroxidase, *IF* immunofluorescence, *Ms* mouse, *Rb* rabbit, *WB* Western blot
